# Flap dynamics in pepsin-like aspartic proteases: a computational perspective using Plasmepsin-II and BACE-1 as model systems

**DOI:** 10.1101/2020.04.27.062539

**Authors:** Soumendranath Bhakat, Pär Söderhjelm

## Abstract

The flexibility of a *β* hairpin structure known as the *flap* plays a key role in catalytic activity and substrate intake in pepsin-like aspartic proteases. Most of these enzymes share structural and sequential similarity. In the apo form of the proteases, a conserved tyrosine residue in the flap region is in dynamic equilibrium between the normal and flipped states through rotation of the *χ*_1_ and *χ*_2_ angles. In this study, we have used apo Plm-II and BACE-1 as model systems. Independent MD simulations of Plm-II and BACE-1 remained stuck either in the normal or flipped state. Metadynamics simulations using side-chain torsion angles (*χ*_1_ and *χ*_2_ of tyrosine) as collective variables sampled the transition between the normal and flipped states. Qualitatively, the two states were predicted to be equally populated. The normal and flipped states were stabilised by H-bond interactions to a tryptophan residue and to the catalytic aspartate, respectively. Further, mutation of tyrosine to an amino-acid with smaller side-chain, such as alanine, reduced the flexibility of the flap and resulted in a flap collapse (flap loses flexibility and remains stuck in a particular state). This is in accordance with previous experimental studies, which showed that mutation to alanine resulted in loss of activity in pepsin-like aspartic proteases. Our results suggest that the rotation of the tyrosine side-chain is the key movement that governs flap dynamics and opening of the binding pocket in most pepsin-like aspartic proteases.

## Introduction

Aspartic proteases play an important role in the life cycle of different pathogens and therefore act as a promising target in structure-based drug discovery. One of the main structural features of aspartic proteases is the presence of the *β*-hairpin conformation often termed as the *flap*.^1^ The flap is a highly flexible part that plays a critical role in the ligand binding by displaying a scissor-like motion necessary for ligand uptake (Figure 1). The conformational flexibility of aspartic proteases and the role of the flap in ligand binding have been studied in several systems, e.g. HIV proteases, ^2,3^ cathepsin-D^4^ and BACE-1.^5–7^ Pepsin-like aspartic proteases^8^ are a subset of aspartic proteases which has a conserved *flap* and *coil* region (Figure 1). The flap acts like a lid which covers the active site of the enzyme. Tyrosine (Tyr) is a conserved residue present in the *flap* of most of the pepsin-like aspartic proteases e.g. plasmepsin I, II, IV, human cathepsin-D, cathepsin-E, BACE-1, BACE-2. It is believed that the rotation of Tyr side-chain plays an important role in the flap dynamics of these enzymes.

**Figure 1:**
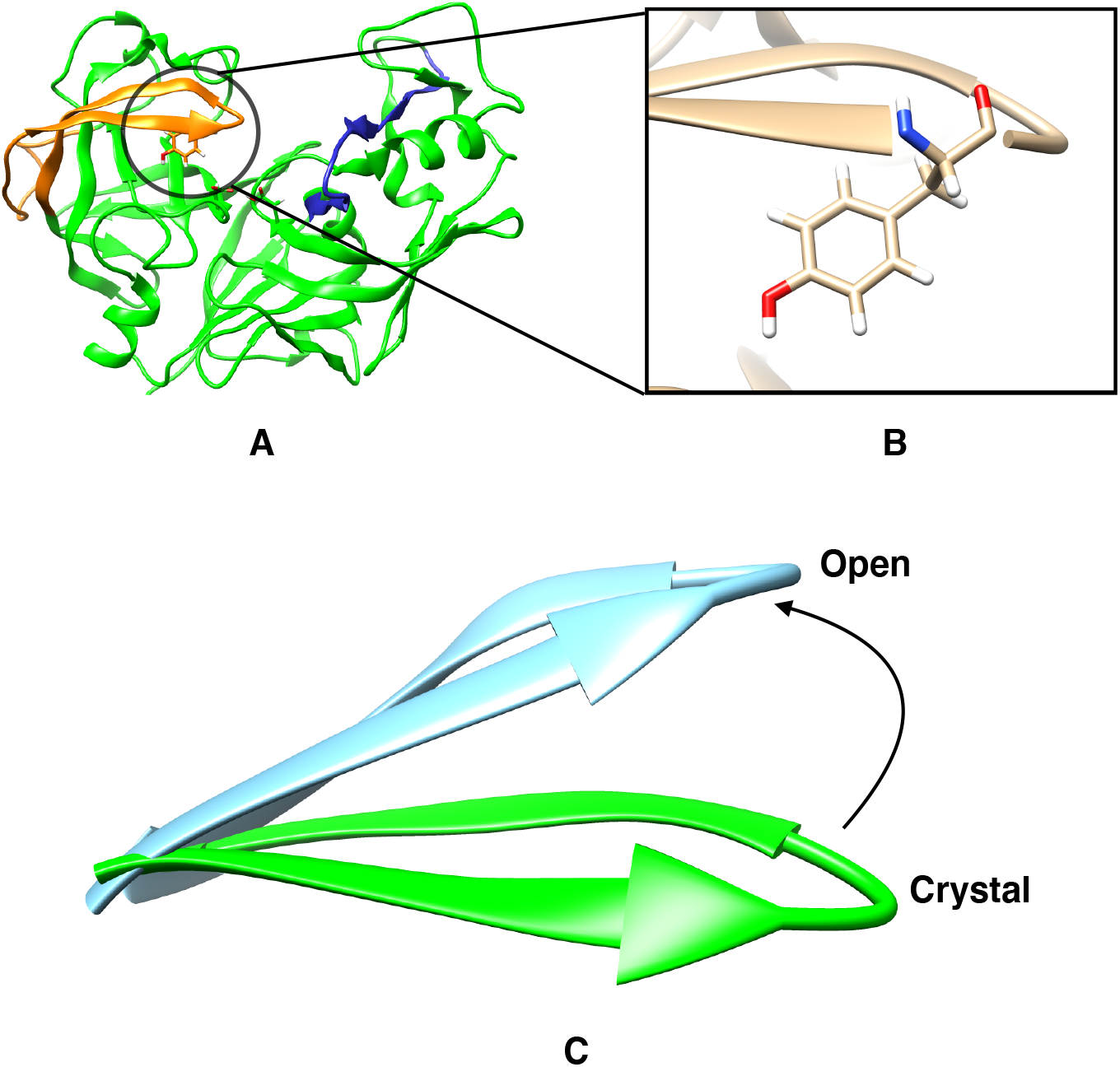
Flap (**Orange**) and coil (**Blue**) region of pepsin-like aspartic protease and location of the conserved tyrosine (Tyr) present in the flap (B). Difference in flap conformation in *crystal* and *open* state (C).

Plasmepsins (*Plm*) are a group of pepsin-like aspartic proteases present in *P. falciparum* (one of the parasites which cause Malaria) and expressed by ten different genes: *Plm* I, II, III, IV, V, VI, VII, IX, X and HAP. Due to the *neglected* status of malaria, the conformational flexibility and structural biology of Plm has not generated much attention among structural and computational biologists when compared with HIV protease. ^2,3,9,10^ Inspired by the idea that the flap of HIV-protease spontaneously opens and closes in MD simulation, ^3^ *Karubiu et al*.^11^ used MD simulations and devised relevant distance parameters to understand the flap opening in Plm II. Further, *McGillewie et al*.^12,13^ used the same distance parameters to understand the flap dynamics and conformational flexibility in Plm I, III, IV and V. *Friedman and Caflisch* ^14^ studied the flexibility of Plm II using MD simulations and hinted that the conserved tyrosine (Tyr) residue present in the flap region might play a role in flap flexibility. *Spronk and Carlson* ^7^ performed classical MD simulations on a homologous protein, BACE-1, and predicted that the side-chain configuration of Tyr governs the conformational flexibility of the flap. They reported three different configurations involving H-bond interactions between Tyr and three neighbouring amino acids (Trp, Ser and Asp). MD simulations sampled a *self-inhibited* conformation, ^7,15^ where the Tyr points toward the catalytic region and forms hydrogen bonds with the catalytic aspartate. The presence of this orientation is not experimentally reported in any BACE-1 or Plm-II crystal structures to date. Due to lack of transitions between these conformations in MD simulations, it was difficult to estimate their free energy differences.

Enhanced sampling methods such as metadynamics, ^16–18^ replica-exchange molecular dynamics (REMD),^19^ umbrella sampling, ^20^ conformational flooding ^21^ etc, have been used regularly to overcome the sampling problem associated with MD simulation. In particular, metadynamics,^16–18^ which uses a repulsive bias potential along some reaction co-ordinates, also known as collective variables (CVs). The addition of a bias potential pushes the system away from local free-energy minima and accelerates sampling of the conformational space.^22^

The aim of our study is to use MD and metadynamics simulations to understand the conformational dynamics in pepsin-like aspartic proteases. Most of these proteases possess structural and sequence similarity. Hence, understanding the conformational dynamics associated with a few of these enzymes can help us understand other homologous enzymes. We use two homologous proteases, Plm-II (Plasmepsin-II) and BACE-1 (*β* secretase-1) as models to investigate 1) what is the role of the Tyr residue in the flap dynamics and 2) whether the rotation of *χ*_1_ and *χ*_2_ torsional angles of Tyr dictates different flap conformations? We test several different collective variables within the metadynamics framework and discuss their effectiveness in sampling flap conformations. We also test the influence of force fields and water models on the population of flap conformations, and suggest possible directions of future work.

## Computational Methods

### Starting structure: Plm II

The co-ordinates of the apo Plm II (PDB ID: *1LF4*)^23^ was retrieved from protein data bank. Prior to simulation, the protonation state of the structure was adjusted using the *H++* server^24^ and missing cysteine–cysteine sulphur–sulphur (S-S) bridges were created. The catalytic aspartic acid at position 214 was protonated while the other one (Asp34) remained negatively charged.

### Starting structure: BACE-1

The co-ordinates of the apo BACE-1 were retrieved from PDB (PDB: 1W50 (SO), 3TPL (SN) and 1SGZ (SSO)). These structures differs in terms of orientation of tyrosine and extent of flap opening (Figure S1 in Supplementary Informations). The structures of BACE-1 were prepared similarly that of Plm-II.

### MD simulations

#### TIP3P water model

The protein was represented by the FF14SB^25^ force field and immersed into a truncated octahedron box with TIP3P^26^ water molecules. The system was neutralised by 9 sodium ions. The box was set such that no protein atom was within 1.0 nm of the box edge. Initial restrained minimisation was performed with steepest descent algorithm for 750 steps followed by 1750 steps of conjugate gradients with a restrained potential of 40 kJ/mol applied on the *C*_*α*_ atoms. The restrained minimisation was followed by 200 steps of unrestrained minimisation. The minimisation was followed by gradual heating (from 0K to 300K) for 400 ps with a harmonic restrained of 40 kJ/mol applied on the *C*_*α*_ atoms and a Langevin thermostat with a collision frequency of 1*ps*^*−*1^ using NVT ensemble. The system was subsequently equilibrated at 300K in an NPT ensemble for 5 ns without restraint and a Berendsen barostat^27^ used to maintain the pressure at 1 bar. Long-range electrostatics were treated using particle mesh Ewald (PME) method^28^ with a grid spacing of 0.1 nm and van-der Waals (vdW) cut-off of 1.2 nm. Finally, independent production runs (using apo Plm-II and BACE-1, Table S1 and S2 in Supplementary Informations) with time step of 2 fs were performed with different initial starting velocities and LINCS algorithm ^29^ was used to constrain the bonds of all hydrogen atoms during the simulation.

#### TIP4P-Ew water model

We have also carried out independent MD simulations using the TIP4P-Ew^30^ water model (referred to as TIP4P-Ew from here on). Simulation parameters were identical as in the TIP4P-Ew simulations. The system was equilibrated in the NPT ensemble using the Berendsen barostat for at least 5 ns. Independent production runs (using apo Plm-II, Table S1 in Supplementary Informations) were performed with time step of 2 fs.

### Metadynamics and Choice of CVs

In metadynamics simulation,^16^ an external bias potential is constructed in the space of a few selected degrees of freedom, known as collective variables (CVs). This potential is built as a sum of Gaussians deposited along the trajectory in the CVs space. The applied bias potential in metadynamics pushes the system away from a local energy minimum into new regions of phase space. In standard metadynamics simulation, gaussians of a predefined height are added during the course of metadynamics. As a result, the system is eventually pushed to explore high free-energy regions. This problem is solved by applying the well-tempered metadynamics protocol,^31^ where the height of the gaussian is decreased with simulation time which eventually leads to smooth convergence in a long time scale. We have used principal component analysis (PCA), time-independent component analysis (TICA),^32^ distances (COM) and torsion angles (Torsion-MetaD) as CVs during our metadynamics investigations.

### Torsion Angles

We have used *χ*_1_ and *χ*_2_ angles of the Tyr residue in the flap (Tyr-77 in case of Plm-II and Tyr-71 in case of BACE-1) as CVs to perform well-tempered metadynamics simulations. All metadynamics simulations were performed at 300K with a gaussian height of 1.2 kJ/mol and a width of 0.05 *radian* deposited every 1 ps. The bias factor was set to 15. Three independent metadynamics simulations (two with TIP3P and one with TIP4P-Ew water model) were performed in case of apo Plm-II. In case of BACE-1, three different metadynamics simulations (using TIP3P water model) were performed with the same settings but starting from the SO, SN and SSO conformations, respectively. For Plm-II, we have also performed metadynamics simulations with CHARMM36 force-field^33^ (with TIP3P water model) using torsion angles as CVs.

### Distance

We have used the distance between the centre of mass (COM) of the catalytic aspartic residues (taking into account only *C*_*α*_ atoms of Asp34 and Ash214) and the COM of the flap region (taking into account all *C*_*α*_ atoms of residues 58-88) as the first collective variable (CV1). The distance between the COM of the catalytic aspartic residues and the COM of the coil region (taking into account all *C*_*α*_ atoms of residues 282-302) was used as the second collective variable (CV2) (Figure S2 in Supplementary Informations). CV1 and CV2 were used in a 2D WT-MetaD simulation with the TIP3P water model. Metadynamics simulations were performed at 300K with a gaussian height of 1.2 kJ/mol and a width of 0.02 nm deposited every 20 ps, and a bias factor of 10. An upper and lower wall at 2.05 nm and 1.60 nm respectively, were placed along CV1, whereas in case of CV2, the upper and the lower wall were placed at 1.75 nm and 1.35 nm, respectively.

### Principal Components

Principal component analysis (PCA) was performed in order to capture the most prominent motions from our MD simulations.^34^ Trajectories from two independent MD runs in TIP4P-Ew water model (starting with apo Plm-II) were combined and PCA analysis was performed on *C*_*α*_ atoms of the protein (ignoring the tail part, Gly0–Asn3) using the *g covar* tool integrated with *Gromacs*. We have used the first two eigenvectors (PC1 and PC2) as CVs in a 2D WT-MetaD with the TIP3P water model. The temperature was set at 300K with a Gaussian height and width of 1.0 kJ/mol and 0.00025 nm, respectively. The bias factor was set at 10. Walls were applied along the eigenvectors to restrict the sampling of regions of high free energy.

### Time-independent component analysis

Time-independent component analysis^1^ was performed using *MSM Builder 3*.*8* ^35^ using one representative MD simulation with TIP4P-Ew water model. Rotational and translational degrees of freedoms were removed from the MD trajectory before performing the dimensionality reduction. The dimensionality reduction started by transforming raw cartesian co-ordinates into a subset of important dihedral features using *DihedralFeaturiser*. We used *χ*_1_ and *χ*_2_ angles associated with the flap region (residues 73-83) of apo Plm II to generate the features (Figure S4 in Supplementary Informations). High-dimensional dihedral features were then reduced to 10 time-lagged independent components by performing TICA using a *lag time* of 10. The TICA features were transferred to a *Plumed* input file via a *Python* script. We were mainly focused on the first 5 TICs. *Plumed Driver* was used to extract the projection of the first 5 TICs from MD trajectories. Finally, well-tempered metadynamics was performed using TIC1 and TIC3 (Figure S3) as CVs with a Gaussian width and height of 0.06 and 1.2 kJ/mol at 300K. The bias factor was set at 15.

## Results

### Flipping of the tyrosine side-chain

Based on earlier investigations, we expected the conserved tyrosine residue in the flap (Tyr-77 in Plm-II; Tyr-71 in BACE-1) to play important roles in the dynamics of the flap. Indeed, MD simulations of apo Plm-II and BACE-1 showed that the hydroxyl group of this residue was interchangeably involved in several hydrogen-bond interactions, and that *slow* conformational dynamics occurred around this residue, on a timescale of at least hundreds of nanoseconds. However, a closer analysis of the dynamics revealed that each particular hydrogen bond was not exceptionally strong. Instead, the high kinetic barriers appeared to occur between the rotational states of the *χ*_1_ torsional angle of the tyrosine side-chain, with each state being able to accommodate several different hydrogen bonds. Thus, this torsional angle, possibly together with its neighbour *χ*_2_, should be a good candidate for biasing in enhanced-sampling simulations.

Our notation for the *χ*_1_ and *χ*_2_ angles of the tyrosine residue is presented in Figure 2.

**Figure 2:**
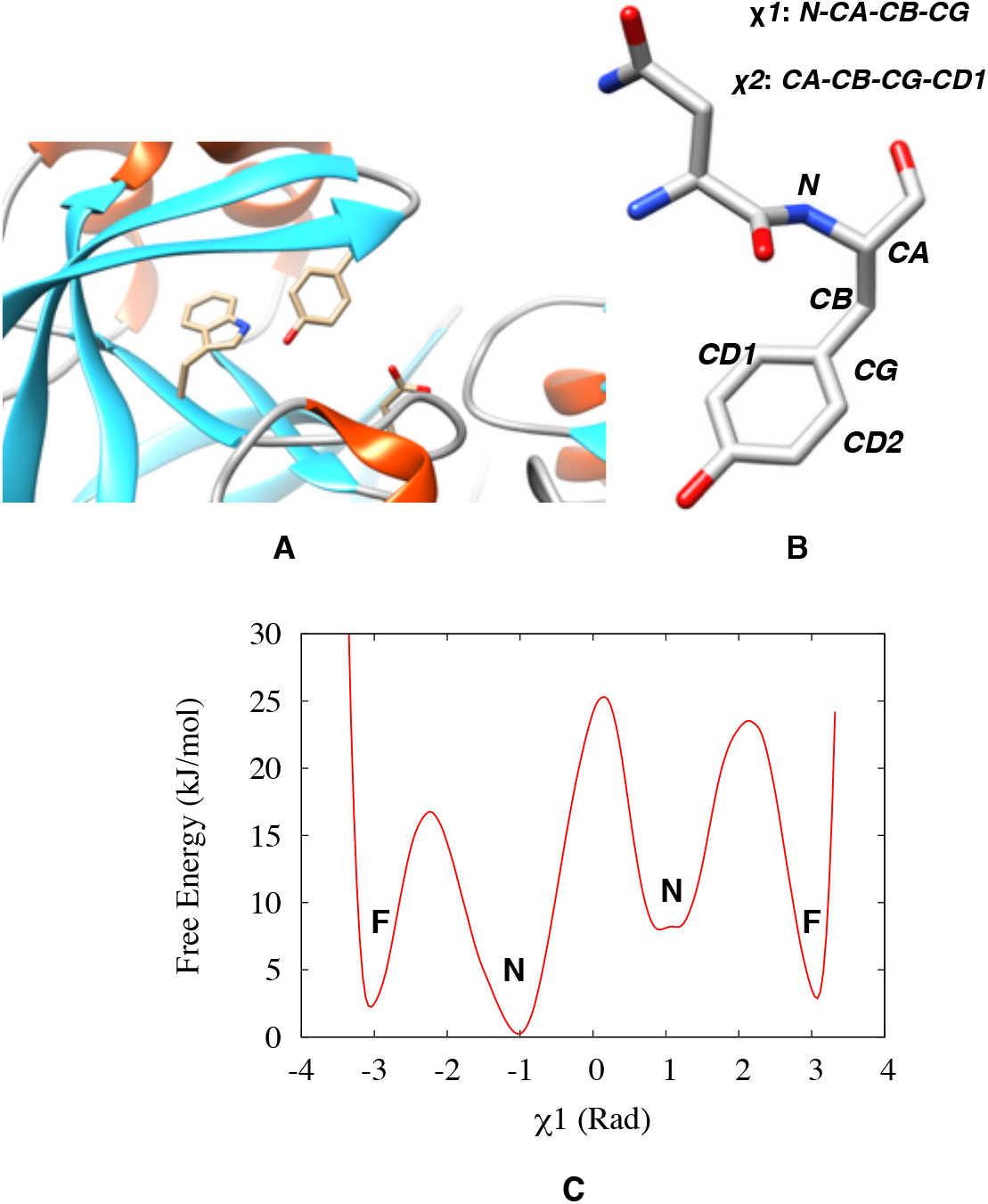
Location of the Tyr,Trp,Asp triad in a typical pepsin-like aspartic protease, Plm-II (A). Definition of the *χ*_1_ and *χ*_2_ angles of Tyr (B). Typical distribution of the *χ*_1_ angle (C), where **N** and **F** denote the normal and flipped states, respectively.

The distributions of the *χ*_1_ angle centred around 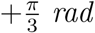 or 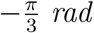 are denoted as *normal*, whereas the distribution centred around ±*π radian* is denoted as *flipped* (Figure 2). Crystal structures of apo Plm-II are always in the normal state, whereas crystal structures of apo BACE-1 (Figure S1 in Supplementary Information) have captured the residue in different states.

### Plasmepsin-II

During MD simulations of apo Plm-II, Tyr-77 formed five major H-bond interactions: with Trp-41, Ser-37, Asp-34, Asn-39 and Gly-216, respectively (Figure 3). The distribution of the *χ*_1_ angle of Tyr-77 showed that only one of the four independent MD simulations^2^ sampled both the normal and flipped states (Figure 4). The normal state was stabilised by formation of interchanging H-bonds to Trp-41, Ser-37 and Asn-39, whereas the flipped state was stabilised by H-bonds to Asp-34 (Figure S12 in the Supplementary Information). Both these states also sampled the *solvent-exposed* conformations, in which the Tyr-77 sidechain did not form any H-bond interactions with neighbouring residues but only with the solvent. Lack of sampling and a great variation of population among the independent MD runs prevented us from predicting the free energy difference between different conformational states of Tyr.

**Figure 3:**
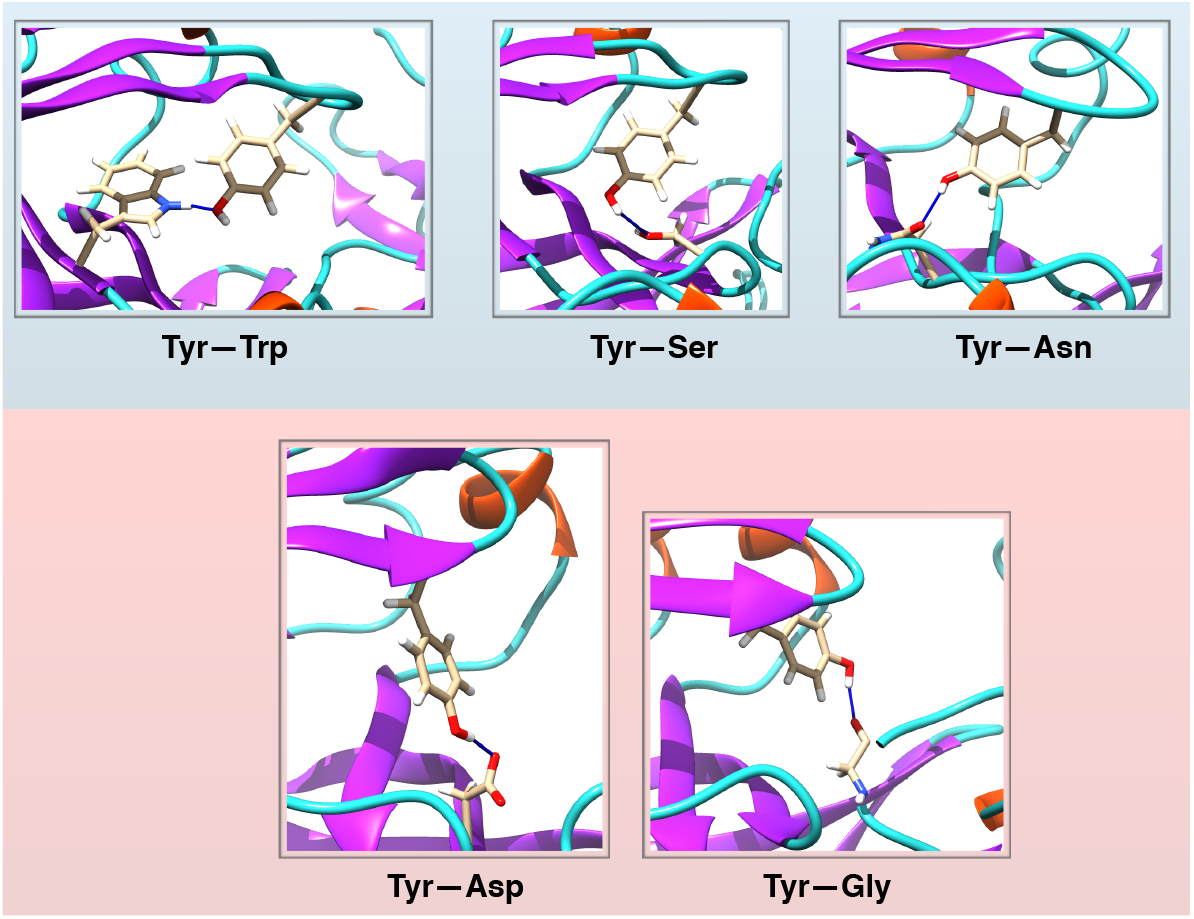
H-bond interactions involving Tyr-77 in Plm-II, divided into those typical for the normal state (upper panel) and flipped state (lower panel).

**Figure 4:**
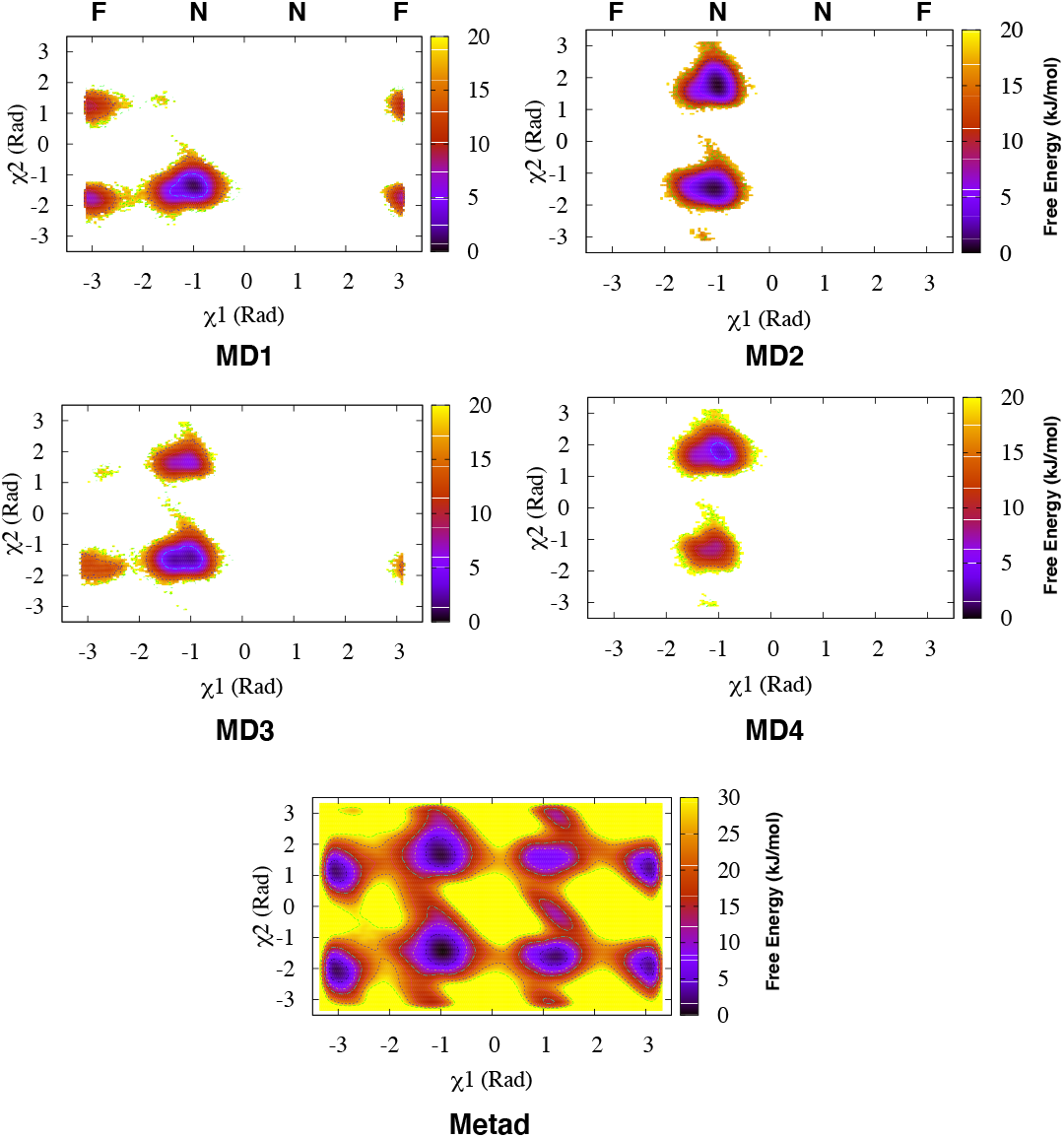
Apparent free-energy surfaces (with respect to the Tyr-77 torsions) for each of the four independent MD simulations of Plm-II (denoted as MD **1-4**) with the TIP3P water model, and from the combined Torsion-Metad simulations (denoted as Metad). The normal (**N**) and flipped (**F**) states are marked on the *χ*_1_ axis.

Metadynamics simulations with biasing on the *χ*_1_ and *χ*_2_ angles (hereafter denoted Torsion-Metad) enhanced the sampling of the rotational degrees of freedom of the Tyr sidechain, which led to an enhanced transition between the normal and flipped states (Figure 4). Qualitatively, the flipped and normal states were found to be equally populated (see Table S6 in Supplementary Informations) with a barrier of more than 10 kJ/mol. Similarly to MD, the metadynamics simulations sampled several interchanging H-bond interactions involving Tyr-77, and the sampling of different H-bonds was significantly better than in MD (Figure 5). The normal state was again stabilised by interchanging H-bond interactions involving Trp-41, Asn-39 and Ser-37, whereas the flipped state was stabilised by H-bonds to Asp-34 and Gly-216 (Figure 5 and Figure S14 in the Supplementary Information). The H-bonds to Trp-41 and Asp-34 were found to be the dominant interactions stabilising the normal and flipped states, respectively (Figure 5).

**Figure 5:**
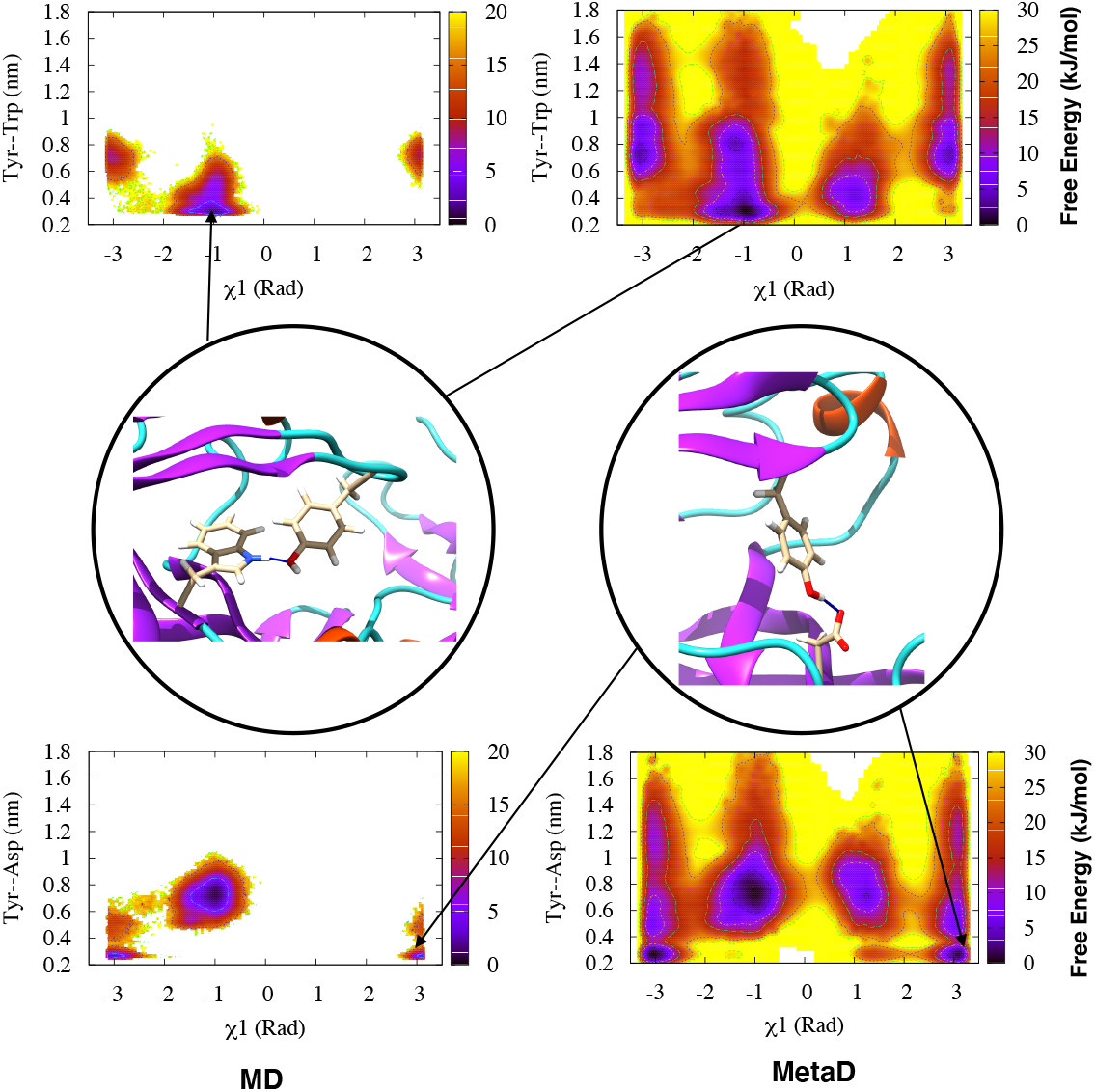
Free-energy surfaces for Plm-II reweighted on *χ*_1_ and H-bond distances to Trp-41 (upper panel) and Asp-34 (lower panel), for the MD (four runs combined) and Torsion-Metad simulations, respectively. Typical structures corresponding to the two H-bonds are also shown.

### BACE-1

MD simulations of BACE-1 were initialised using three different crystal structures, denoted SO, SN, and SSO. These structures differ in terms of orientation of Tyr-71 and extent of flap opening (Figure S1 in the Supplementary Information). The MD simulations starting from the SO and SN crystal structures only sampled the normal state (Figure 6) of Tyr-71, which was stabilised by H-bond interaction to Trp-76 (comparable to Trp-41 in apo Plm-II). On the other hand, MD simulations started from the SSO crystal structure remained stuck in the flipped state (Figure 6), which was stabilised by formation of a H-bond to Asp-32 (comparable to Asp-34 in Plm-II) (Figure S15 in the Supplementary Information).

**Figure 6:**
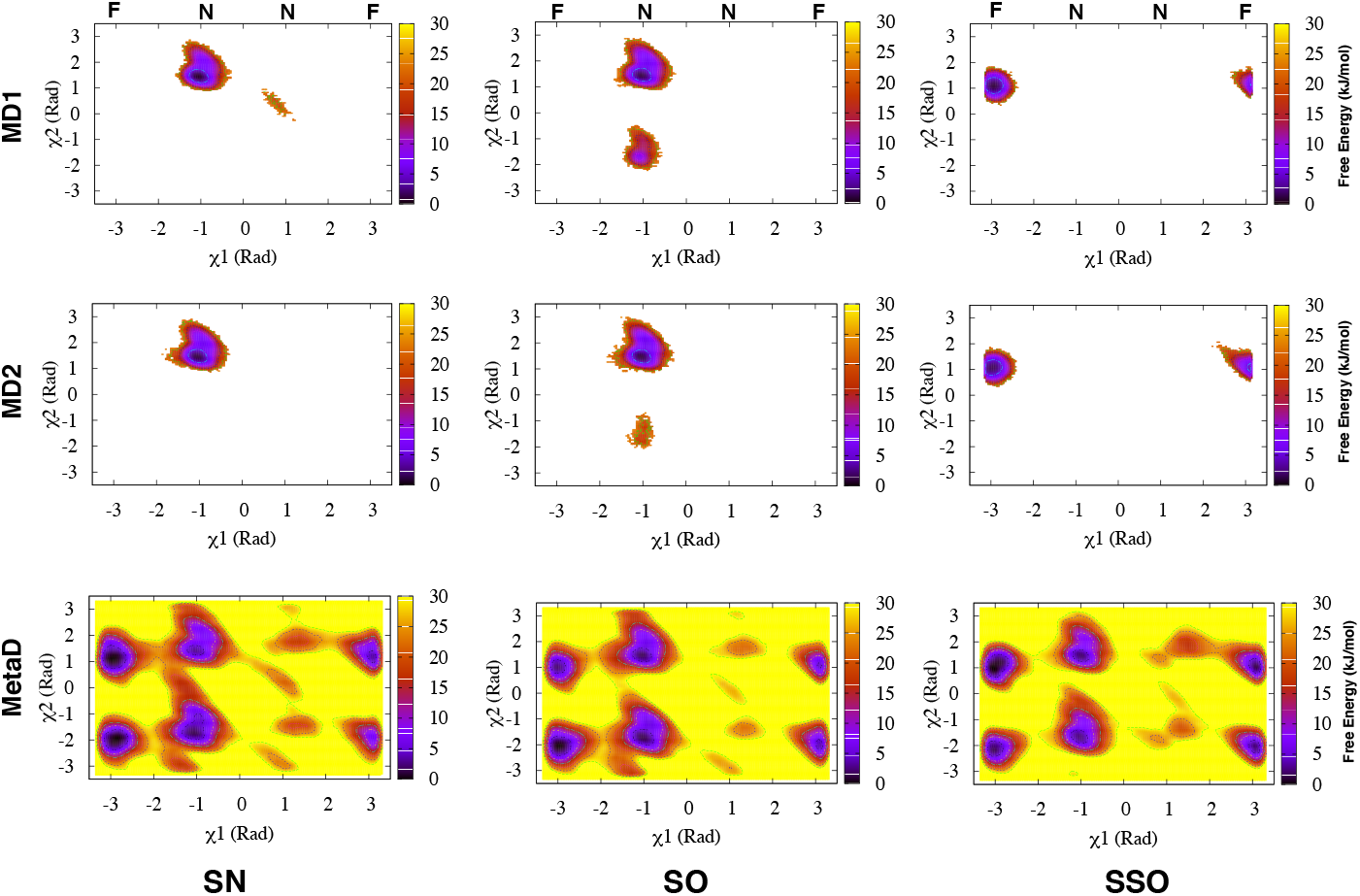
Apparent free-energy surfaces of BACE-1 from MD and Torsion-Metad simulations starting from the SN, SO and SSO structures, respectively. Independent MD simulations starting from the SN or SO structures only sampled the normal (**N**) state (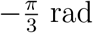, whereas simulations starting from the SSO structure only sampled the flipped (**F**) state. Torsion-Metad simulations starting from the SN, SO or SSO structures sampled both the **N** and **F** states.

Torsion-Metad simulations enhanced transitions between the normal and flipped states (Figure S8 and Figure S15 in the Supplementary Information). The sampling of these two states in BACE-1 is comparable with Plm-II. H-bond interactions to Trp-76 and Asp-32 dominate the normal and flipped states, respectively (Figure 7). The metadynamics simulations also sampled H-bonds to Ser-35 and Lys-107. The H-bond to Lys is caused by an unusual backwards orientation of Tyr-71, which we define as the *Tyr-back* conformation. The free-energy difference between the H-bonds to Trp-76 and Ser-35 was predicted to be ∼ 7 kJ/mol. The difference in free energy between the H-bonds to Trp-76 and Lys-107 was ∼ 20 kJ/mol (Figure 7).

**Figure 7:**
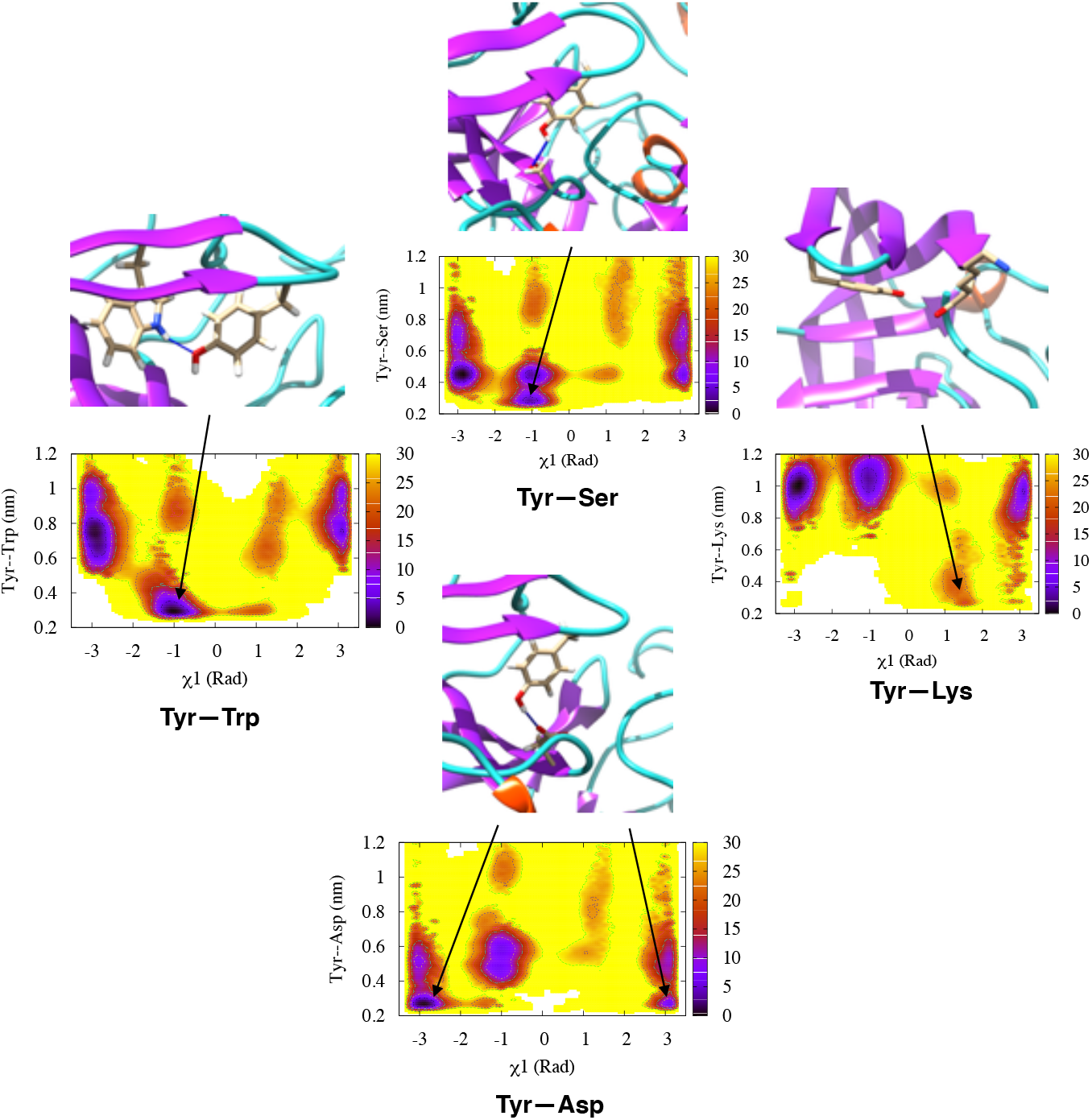
Free-energy surface for the combined Torsion-Metad simulations on BACE-1 reweighted on *χ*_1_ and different H-bond distances (Trp-76, Ser-35, Asp-32 and Lys-107). The corresponding free-energy surfaces for the individual runs (starting with the SN, SO and SSO conformations) can be found in the Supplementary Information.

### Implications for flap opening

#### Plasmepsin-II

To describe large-scale motion of the flap, especially *opening* of the flap, we defined two distances as illustrated in Figure 8. For each MD simulation, the free energy surface was calculated from the corresponding probability distribution with respect to DIST2 and DIST3 as well as to DIST2 and *χ*_1_. The lack of correlation between DIST2 and DIST3 indicates that the flap and the coil regions move largely independently. The extent of flap opening (DIST2) and the rotation of the *χ*_1_ angle of Tyr-77 are clearly related. When Tyr-77 is in the normal state, a broad basin around DIST2≈ 1.2 nm is stabilised by interchanging H-bonds to Trp-41, Ser-37 and Asn-39. In the additional basin around DIST2≈ 1.6 nm seen in some simulations, there are solvent exposed conformations of Tyr-77 but also conformations with H-bond to Trp-43. When Tyr-77 is in the flipped state, the free-energy minimum centred around DIST2≈ 1.05 nm is stabilised by interchanging H-bonds to Asp-34 and Gly-216. Finally, the flap adapts *open* conformations around DIST2 2.0 nm (conformational snapshots corresponding open conformations are highlighted in Figure S20 in Supplementary Information).

**Figure 8:**
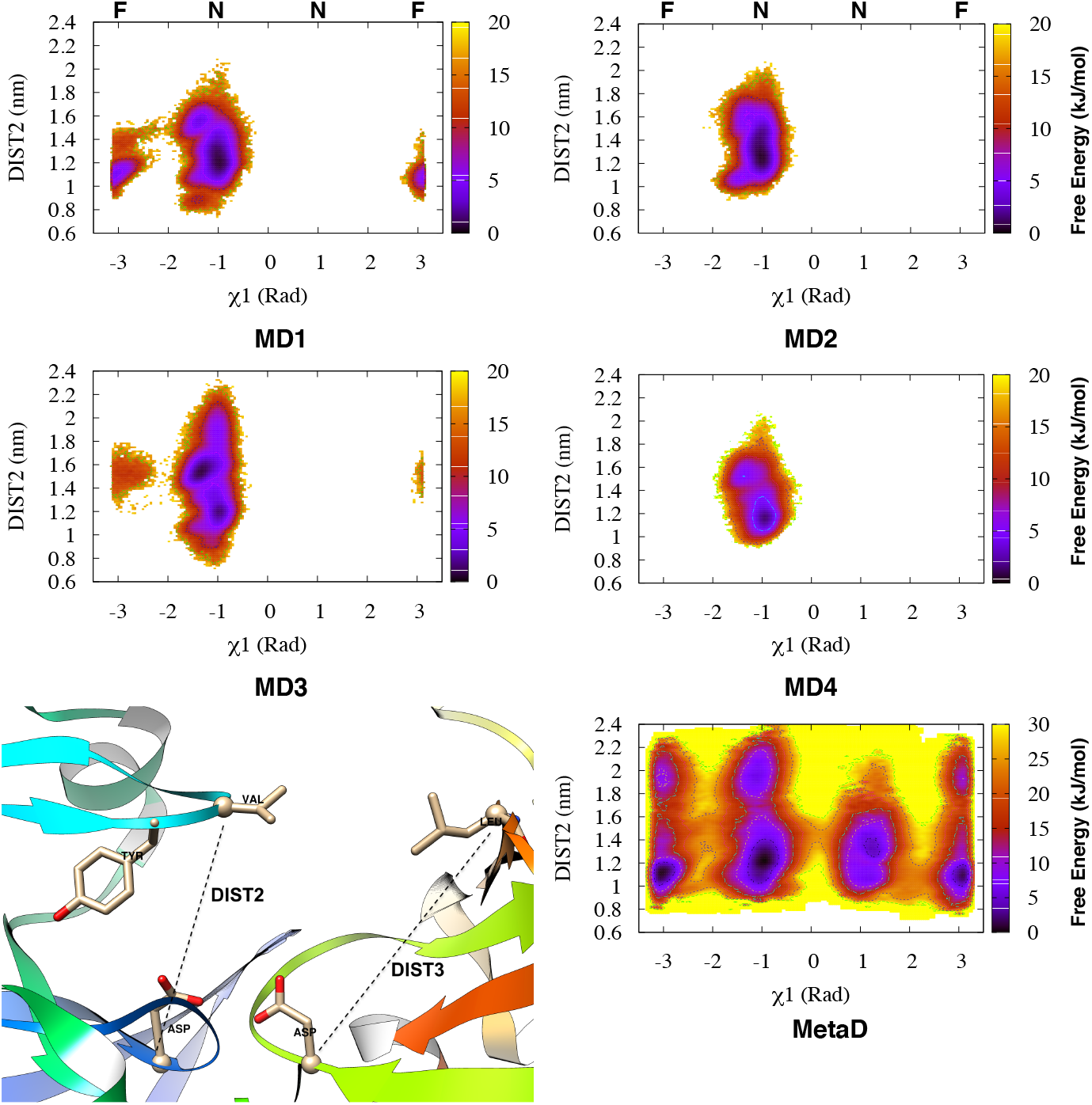
Apparent free-energy surfaces for Plm-II reweighted on *χ*_1_ and DIST2, showing sampling of overall flap flexibility during MD (denoted as MD**1-4**) and Torsion-Metad simulations. **N** and **F** denote the normal and flipped states of Tyr-77, respectively.

All four independent MD simulations with TIP3P water model sampled free-energy minima with DIST2 (Figure 8) centred around ∼1.2 nm and ∼1.6 nm (Figure S21 and Figure S22 in Supplementary Information shows conformational snapshots corresponding each basin) respectively. In MD simulations, the free-energy basin around ∼1.2 nm sampled only interchanging H-bonds to Trp-41 and Ser-37. Only one of the four independent simulations sampled around ∼1.05 nm (stabilised by the H-bond to Asp-34). Moreover, one of the four independent MD simulations sampled the flap opening. More structural details can be found in the clustering section of the Supplementary Information.

Torsion-Metad simulations gave a better picture of the flap dynamics in Plm-II (Figure 8). Metadynamics simulations sampled typical minima with DIST2 centred around ∼1.05 nm, ∼1.2 nm and ∼1.6 nm. Free-energy minima centred around ∼1.05 nm and ∼1.2 nm were found to be equally populated^3^ and the free energy cost of flap opening (transition from a DIST2 of ∼ 1.05 nm to ∼ 2.0 nm) was predicted to be ∼ 3 kJ/mol^4^.

The sampling of the open conformation was not directly influenced by the enhanced sampling of the Tyr-77 side-chains (Table S10 in the Supplementary Information). The only significance appears to be that one avoids being trapped in the normal or flipped states. Once the H-bond interactions between Tyr-77 and neighbouring residues are broken and it becomes solvent-exposed, the sampling of the flap opening is in fact similar among standard MD simulations and metadynamics simulations.

#### BACE-1

We have devised analogous DIST2 and DIST3 descriptors for BACE-1 (see Supplementary Information). The flap opening (DIST2) and rotation of the *χ*_1_ angle of Tyr-71 are related (Figure 9) in a similar way as for Plm-II. Thus, when Tyr-71 is in the normal state, the free-energy minimum centred around DIST2≈ 1.2 nm is mainly stabilised by interchanging H-bonds to Trp-76, Ser-35 and Lys-107. When Tyr-71 is in the flipped state, the free-energy basin centred around DIST2≈ 1.05 nm is stabilised by formation of a H-bond to Asp-32.

**Figure 9:**
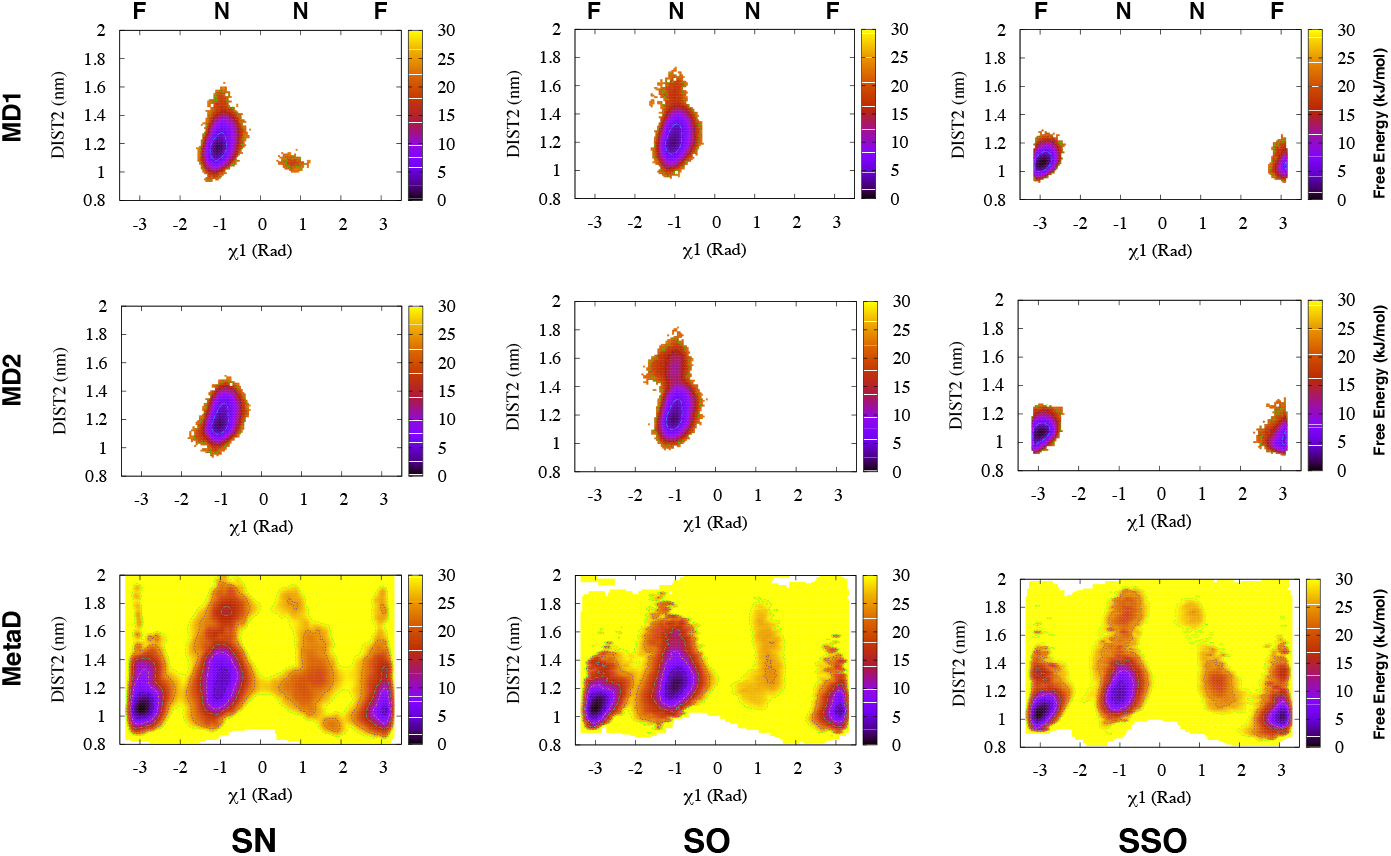
Free energy surface reweighted on *χ*_1_ and DIST2 in MD and Torsion-Metad simulations starting with SN, SO and SSO conformations. **N** and **F** denote the normal and flipped states of Tyr-71.

Figure 9 again shows the incomplete sampling of the MD simulations, whereas the Torsion-Metad simulations sampled both states. The free-energy basins centred around a DIST2 of ∼1.2 nm and ∼1.05 nm, respectively, were found to be equally populated^5^. This suggests that the flap is in dynamic equilibrium between the normal and flipped conformations (Figure 9). The largest flap opening (DIST2≈ 1.8 nm) was observed in the Torsion-Metad simulation started from the SN conformation.

### Extent of opening needed for ligand release

In order to understand the extent of flap opening necessary to accommodate the ligand in the active site, we have performed a well-tempered metadynamics simulation of a *Plm-II–ligand* complex. The distance between the centre of mass of the active site residues and the ligand heavy atoms was selected as CV to accelerate ligand unbinding. This type of simulation will not necessarily follow a low-energy path, but may anyway give a rough picture of the unbinding process. Plotting DIST2 as a function of time (or equivalently the stage of unbinding) indicated that a flap opening (DIST2) of ∼2.0 nm (corresponding to the open conformation in the apo simulations) is sufficient for ligand binding/unbinding (Figure 10). Tyr-77 remained in the normal state during the unbinding event. It is important to note that the amount of flap opening might depend on the size of the ligand. We chose a small molecule inhibitor with hydroxylethylamine scaffold (Figure S18 in Supplementary Information) as the ligand, owing to the importance of such inhibitors in antimalarial drug discovery.^36^

**Figure 10:**
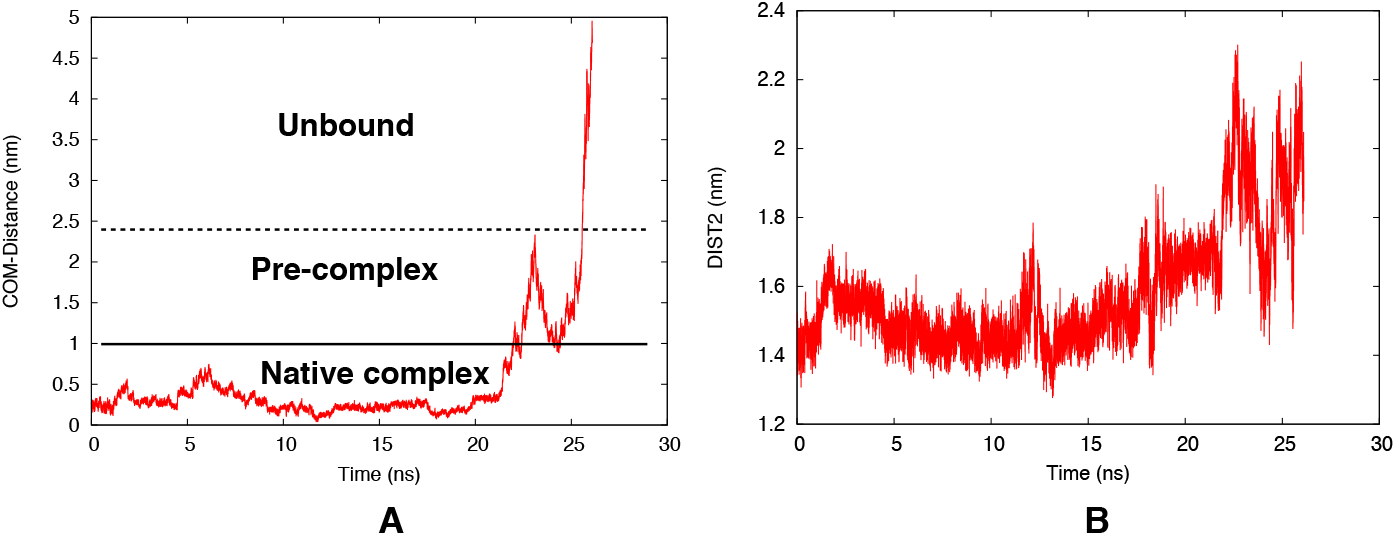
Time evolution of the unbinding CV (distance between the centre of mass of the active site residues and ligand heavy atoms) during the metadynamics simulation (A). Time evolution of DIST2 showing the extent of flap opening during ligand exit (B). The stability of the ligand in active site has been highlighted in Figure S23 in Supplementary Information.

### Forcefield, watermodel and choice of CVs

All results presented up to this point were obtained using the FF14SB force field for the protein and the TIP3P water model. In order to test the effect of the force field and water model on the population of the normal and flipped states, we used apo Plm-II as model system.

### Force field

A Torsion-Metad simulation with the CHARMM36 force field and the TIP3P water model sampled both the normal and flipped conformations of Tyr-77. The flipped state was predicted to be slightly more populated compared to the normal state. The free-energy difference between the flipped and normal state was predicted to be ∼3 kJ/mol (Table S6 in the Supplementary Information).

### Water model

We also carried out four independent MD simulations of apo Plm-II with the TIP4P-Ew water model (and the FF14SB force field). Two of the four independent MD simulations sampled both normal and flipped states (Figure S13 in the Supplementary Information). Besides H-bond interactions to Trp-41, Ser-37, Asn-39 and Asp-34; these TIP4P-Ew simulations sampled an additional H-bond to Gly-216 (Figure S13 in the Supplementary Information).

Torsion-Metad simulations with TIP4P-Ew water model sampled both the normal and flipped states. The population of normal and flipped states as well as different H-bond interactions were comparable to the TIP3P results (Figure S16 and Figure S17 in the Supplementary Information).

### Choice of CVs

We also performed metadynamics simulations of apo Plm-II using three other types of collective variables, namely principle components (PCA), centre-of-mass distances (COM), and time-lagged independent components (TICA), as described in the Supplementary Information). Metadynamics simulations using PCA and COM CVs did a poor sampling of the normal and flipped states (Figure S16 in Supplementary Information). This is due to the fact that the slow degrees of freedom (*χ*_1_ and *χ*_2_ angles of Tyr-77) were not explicitly biased during these simulations. It is also possible that the rather long simulation used for PCA analysis led to CVs that captured motion *between* two structural basins instead of local fluctuations. This factor could contribute to the slow convergence of the metadynamics simulations.

Dimensionality reduction using TICA was able to capture rotational degrees of freedom associated with Tyr-77. The metadynamics simulation using the first and third TICA components as CVs sampled both the normal and flipped states (Figure S16 in Supplementary Information). Qualitatively, the sampling using TICA metadynamics and Torsion-Metad were comparable to eachother.

### Convergence

#### Plasmepsin-II

The two independent Torsion-Metad simulations using the TIP3P water model reached apparent convergence after around 500 ns. To assess the convergence, we have calculated the free-energy difference between the flipped and normal (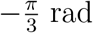) states as a function of simulation time (Figure S7 in Supplementary Information). The last part of each simulation (500 ns onwards) was used for statistical analysis. The average free-energy difference between the flipped and normal states was calculated to be 1.02 ± 0.01 and −0.61 ± 0.01 kJ/mol for the two independent runs (Table S6 in Supplementary Information).

The Torsion-Metad simulations with TIP4P-Ew water model and with the CHARMM force-field, respectively, also reached apparent convergence after around 500 ns (Figure S7 in Supplementary Information). The average free-energy difference between the flipped and normal states was predicted to be 0.62±0.01 kJ/mol and −3.16±0.01 kJ/mol for the TIP4P-Ew and CHARMM simulations, respectively (Table S6 in Supplementary Information).

#### BACE-1

Torsion-Metad simulations starting with SO, SN and SSO conformations reached convergence around ∼300 ns. Statistical analyses were carried out using the last part (300 ns onwards) of the simulation (Figure S8 in Supplementary Informations). The hill heights were very small at this part of simulation. The average free energy difference between flipped and normal state 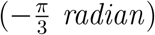 calculated to be −1.68 ± 0.01 kJ/mol, −2.10 ± 0.01 kJ/mol and −3.53±0.02 kJ/mol for SO, SN and SSO respectively (Table S8 Supplementary Information). In the presentations of results in the previous sections, we have only included the Torsion-Metad simulation started from the SN conformation.

### Mutational study

We computationally mutated the conserved Tyr to alanine (Ala) in apo Plm-II and BACE-1. Alanine does not possess a bulky side-chain like Tyr. The apparent free-energy surface on DIST2 and DIST3 shows a complete flap collapse in both Plm-II and BACE-1 (Figure 11 and Figure S19 in Supplementary Information).

**Figure 11:**
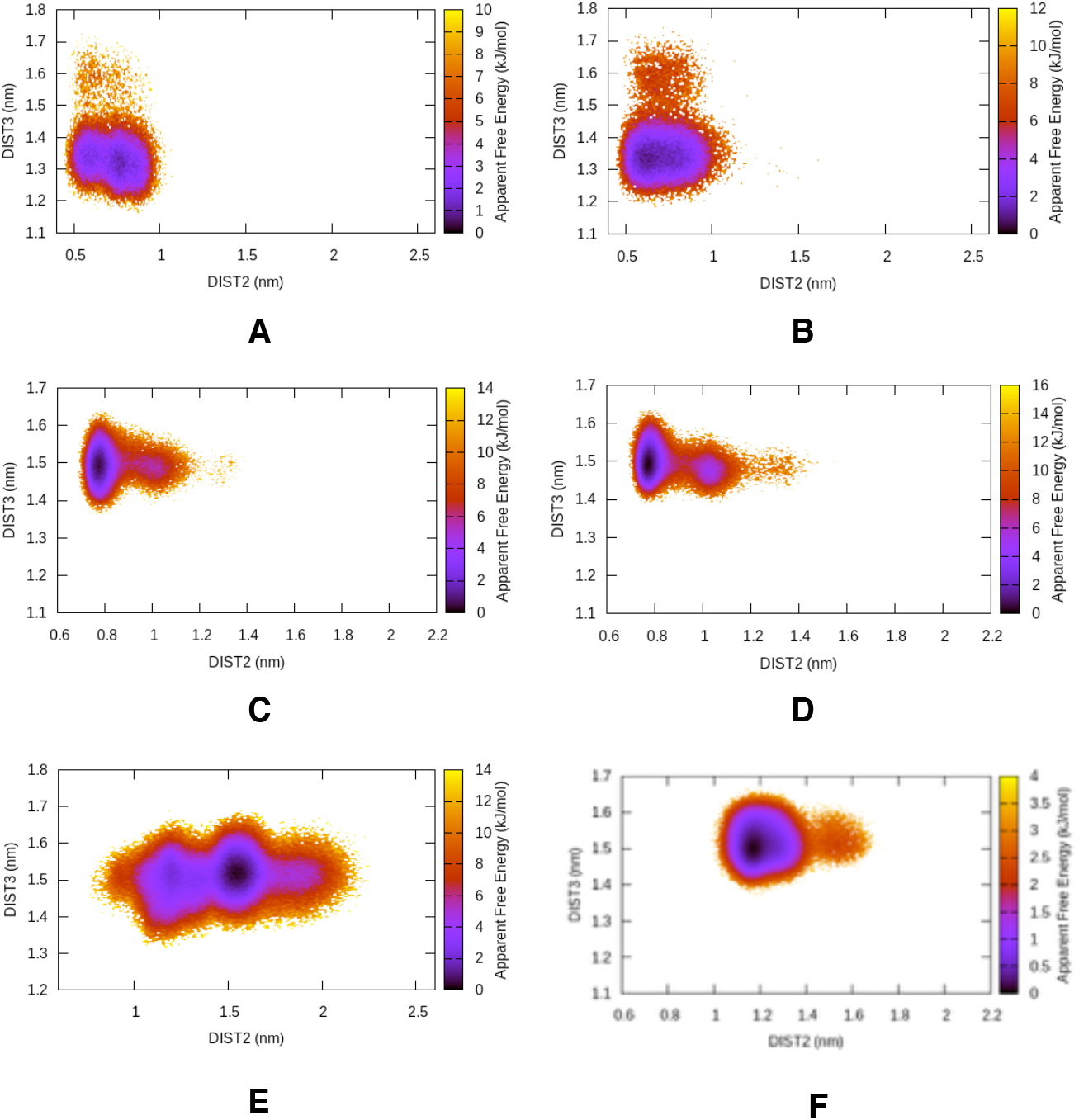
Free-energy surface projected on DIST2 and DIST3 in the case of apo Plm-II (A and B) and BACE-1 (C and D) with Ala mutation. For comparison, the wild type Plm-II and BACE-1 (F) are also shown. Projection on DIST2 highlights that the flap loses its flexibility in the mutant protein.

## Discussion

The flap region of apo Plm-II and BACE-1 adapts different conformations due to rotation of *χ*_1_ and *χ*_2_ angles of the conserved tyrosine residue (Tyr-77 and Tyr-71 in case of Plm-II and BACE-1 respectively). Mutation of Tyr to Ala led to complete flap collapse in Plm-II (Figure S19 in Supplementary Informations) and BACE-1. Previous experimental studies highlighted that mutation of Tyr to Ala resulted in loss of activity.^37^ Mutation of Tyr to other amino acids with smaller side-chains, e.g. Thr, Ile, Val in chymosin (another pepsin-like aspartic protease), also resulted in loss of activity. However, mutation of Tyr to Phe in pepsin didn’t diminish the activity of the protein.^38,39^ Like Tyr, Phe also possesses rotational degrees of freedom along the *χ*_1_ and *χ*_2_ angles. Hence, the flap of pepsin-like proteases with Phe (*Toxoplasma gondii* aspartic proteases, Plm IX, X and Plm V) can still remain dynamic in the apo conformation. ^40,41^

MD simulations sampled both the normal and flipped states. However, a low transition probability between the states makes transitions *rare*. Rotation of the *χ*_2_ angle of Tyr was found to be the slowest degree of freedom separating normal and flipped states. On the other hand, rotation of the *χ*_1_ angle acts as an auxiliary collective variable. Metadynamics simulations with *χ*_1_ and *χ*_2_ angles as CVs sampled the normal and flipped states much more reliably. However, our results do not rule out the possibility that there are other kinetic barriers (enthalpic or entropic) that limit the sampling of flap dynamics in various ways and that are not overcome by our enhanced-sampling approach.

Both the normal and flipped states were stabilised by formation of interchanging H-bonds with neighbouring residues. A common pattern among these interactions is the formation of H-bonds to Trp and Asp (Figure 12). We speculate that the population of normal and flipped states depends on the protonation states of catalytic aspartates. Recent *CpHMD* simulations showed that the population of hydrogen bonds in pepsin-like aspartic proteases depends on the *pH* through the varying protonation of the aspartic dyad.^42^

**Figure 12:**
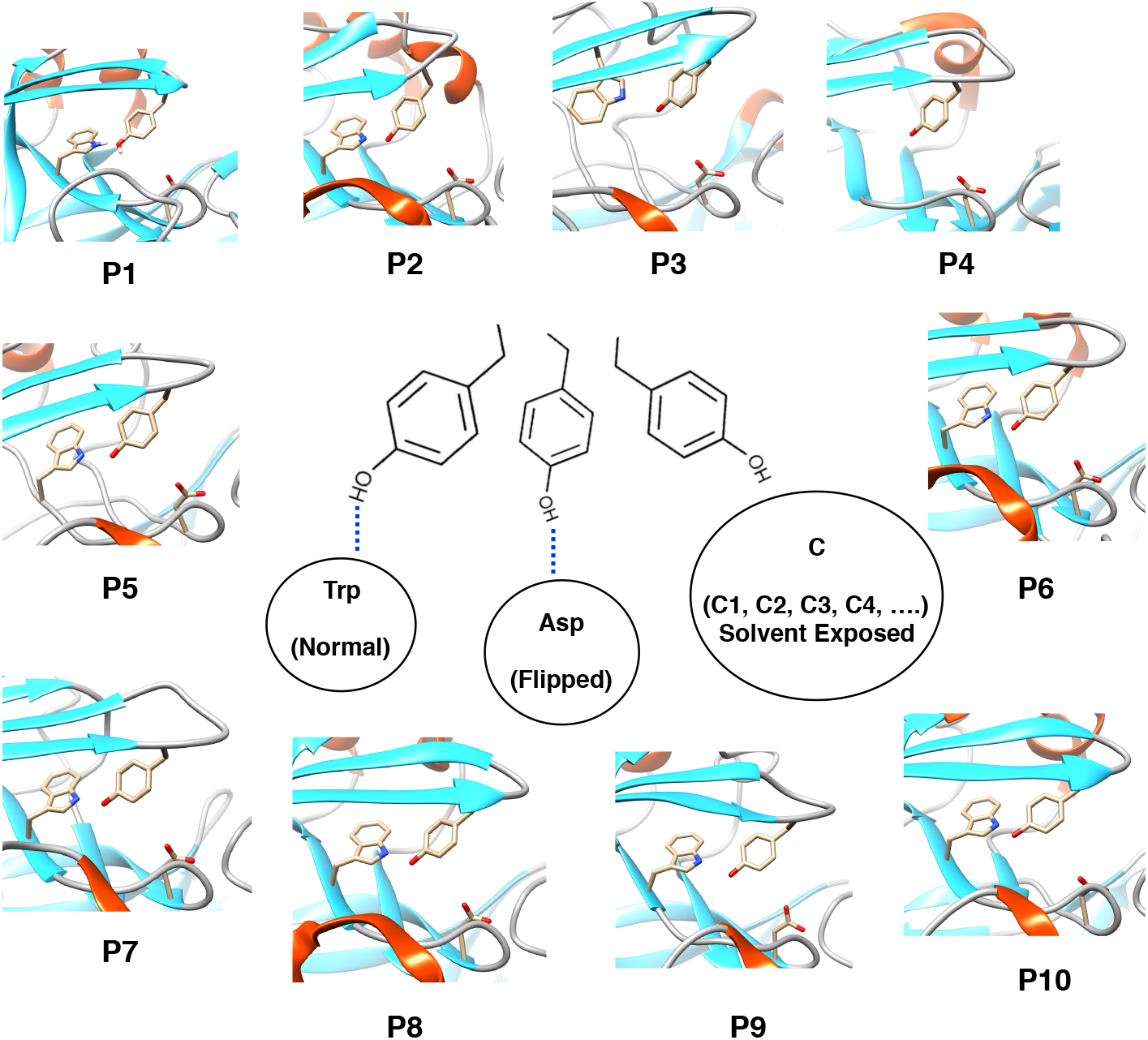
Unified mechanism of conformational dynamics in pepsin-like aspartic proteases: Three different conformational states associated with side-chain flexibility of Tyr in pepsin-like aspartic proteases. The normal and flipped states are typically stabilised by H-bonds to Trp and Asp. C is the solvent-exposed subspace which consists of several rotameric conformations (C1, C2, C3, C4, … etc.) of Tyr. P1 to P10 denotes crystal structures of human cathepsin-D (PDB:1LYA), cathepsin-E (PDB:1TZS), BACE-2 (PDB:3ZKQ), Plm-V (PDB:4ZL4), bovine chymosin (PDB:4AUC), human pepsin (PDB:3UTL), candidapepsin (PDB:2QZW), human renin (PDB:5SY2), Plm-I (PDB:3QS1) and Plm-IV (PDB:1LS5) respectively. All structures (except Plm-V) possessed conserved Tyr, Trp and Asp residues similar to Plm-II and BACE-1. Plm-V is missing the conserved Trp residue.

In *R. pusillus* protease, the normal state is stabilised by a hydrogen bond between Tyr-75 and Trp-39.^43^ In a set of experiments, Park et al. ^44^ replaced Trp-39 with other residues and observed a decreased activity, which suggests a role of Trp in stabilising the normal state of Tyr. The crystal structure of Plm-V from *P. vivax* (PDB: 6C4G,^45^ 4ZL4^46^) does not possess the Trp residue, but the flap region still remains dynamical due to the rotational degrees of freedom associated with Tyr. ^47^

Formation of the H-bond to Asp led to formation of a self-inhibited conformation, which was also observed in previous MD simulations of apo BACE-1.^15^ However, it was not reported in previous MD simulations with apo Plm-II. Crystal structures of Plm-II and bovine chymosin protease show that Tyr can adapt both normal and flipped states (Table 1), which influences the extent of flap opening (Figure 13). These crystal structures gives experimental support to our findings.

**Table 1:** Crystal structures of Plasmepsin-II (PDB: 2BJU, 2IGY, 4Z22) and bovine chymosin protease (PDB: 1CMS) show different conformational states of Tyr. F and N denotes flipped and normal states respectively.

| PDB | $\chi_1$ (radian) | DIST2 (nm) | State |
| --- | --- | --- | --- |
| 2BJU | -3.12 | 1.50 | **F** |
| 2IGY | -2.6 | 1.74 | **F** |
| 4Z22 | -1.02 | 2.11 | **N** |
| 1CMS | 3.08 | 1.28 | **F** |

**Figure 13:**
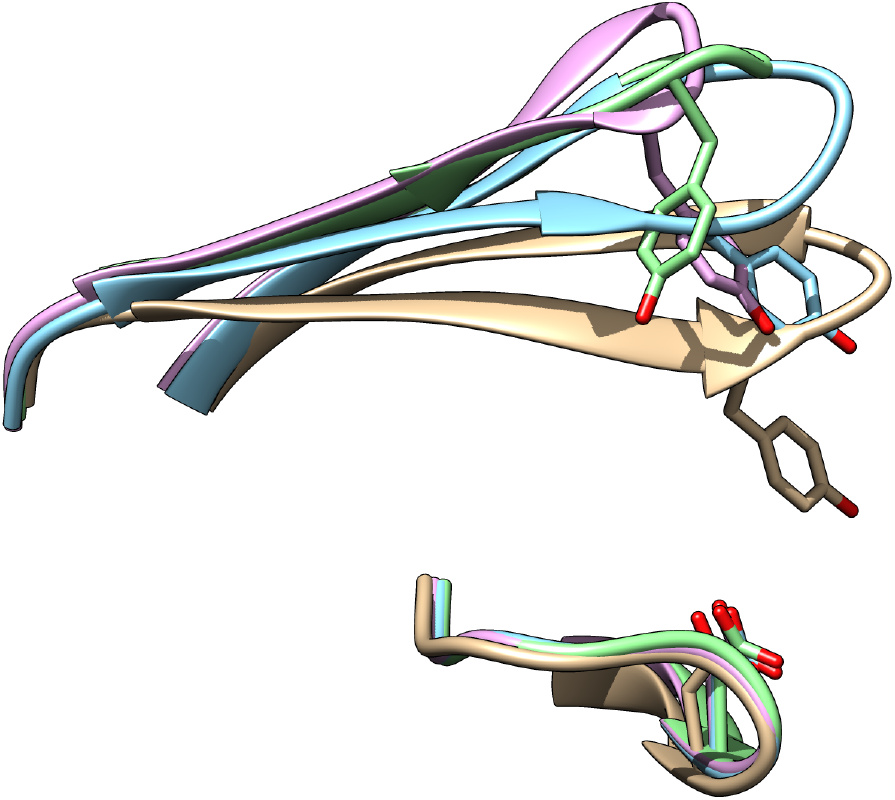
Orientation of tyrosine in crystal structures of Plm-II (PDB: 2BJU (blue), 2IGY (magenta) and 4Z22 (green)) and bovine chymosin protease (PDB: 1CMS (grey)).

Spontaneous flap opening was also sampled during MD and metadynamics simulations. However, the extent of flap opening differed between Plm-II and BACE-1. The metadynamics-based unbinding study provided an overview of the extent of flap opening necessary for substrate entry in Plm-II. In that simulation, Tyr remained in the normal state during ligand unbinding. In the future, more extensive simulation studies are necessary to understand the actual role of Tyr in the substrate binding.

## Conclusion

The dynamic nature of the flap in pepsin-like aspartic proteases is necessary for its catalytic activity and substrate binding. Most of these aspartic proteases possess a conserved triad of Tyr, Trp and Asp (Figure 12). We predict that the flap dynamics in almost all pepsin-like aspartic proteases is governed by the rotational degrees of freedom associated with the Tyr residue. Recently Akke and co-workers^48^ used 13*C* relaxation dispersion experiments to capture kinetics of aromatic side-chain flipping in a small globular protein, BPTI. Furthermore methods such as Markov state model (MSM)^**?**^ and infrequent metadynamics^18^ have been frequently used to predict kinetics associated with conformational transition. We believe that 13*C* relaxation dispersion experiments combined with MSM/infrequent metadynamics can be used in case of pepsin-like aspartic proteases to capture kinetics of tyrosine ring flipping. The orientation of tyrosine directly influences the volume of the binding site (Figure S24 and Figure S25 in Supplementary Information). Drug designers may be able to exploit this property by designing inhibitors of varying size which can lock tyrosine at any desired conformation. We believe that our study will act as a starting point to perform experiments that can validate or modify the structural and mechanistic insights into pepsin-like aspartic proteases.

## Supporting information

Supplementary Informations

## Acknowledgement

PS thanks the Crafoordska Foundation and the Swedish Research Council (project 2017-05318) for financial support. The computations were performed on computer resources provided by the Swedish National Infrastructure for Computing (SNIC) at LUNARC (Lund University) and HPC2N (UmeåUniversity).

## Data and Software Availability

All input files (.tpr, plumed.dat, reweighting protocol, scripts to generate TICA etc.) can be accessed here: https://github.com/sbhakat/Plasmepsin-bace. All simulations were performed using Plumed 2.5 patched with Gromacs 2018.

## Supporting Information Available

Supplementary informations contain computational methods, tables and figures.

## Footnotes

1 using lag time 10

2 Only simulations with the TIP3P water model + FF14SB force-field are considered here, whereas other combinations are discussed in a later section

3 see Table S6 in the Supplementary Information

4 Table S7 in the Supplementary Information

5 see Table S8 in the Supplementary Information

