## Supplementary Informations for "Flap dynamics in pepsin-like aspartic proteases: a computational perspective using Plasmepsin-II and BACE-1 as model systems"


#### Supporting Information Available

#### Computational Methods

Table S1: Length (in ns) of MD simulations starting with TIP3P and TIP4P-Ew water models using apo Plm-II.

| Simulation | Length |
|---|---|
| **TIP3P** | |
| MD1 | 528 |
| MD2 | 562 |
| MD3 | 442 |
| MD4 | 584 |
| **TIP4P-Ew** | |
| MD1 | 669 |
| MD2 | 535 |
| MD3 | 530 |
| MD4 | 530 |

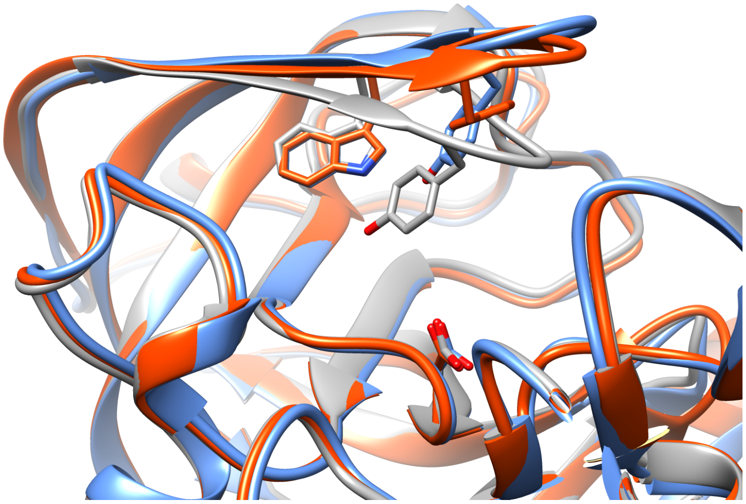

Figure S1: Difference in flap opening and orientation of Tyr in SO (PDB:1W50, *blue*), SN (PDB:3TPL, *grey*) and SSO (PDB:1SGZ, *orange*) conformations of apo BACE-1. In SSO conformation, Tyr points backwards.

Table S2: Length (in ns) of MD simulations starting with SO, SSO and SN conformation of BACE-1. The simulations were carried out in TIP3P water model.

| **Simulation** | **Length** |
|---|---|
| **SO** | |
| MD1 | 774 |
| MD2 | 835 |
| **SSO** | |
| MD1 | 518 |
| MD2 | 513 |
| **SN** | |
| MD1 | 790 |
| MD2 | 791 |

#### Unbinding using metadynamics: Plm-II-ligand complex

Prior to metadynamics simulation on Plm-ligand complex (PDB ID: *4Y6M*),[1] we used the following procedures to prepare the system for simulation. The ligand (Figure S18) was extracted from the complex and was parameterised by the general amber force field (GAFF)[2] whereas the protein was treated using the FF14SB force field. The complex was then neutralised using 5 sodium ions and immersed into a truncated octahedral box with TIP3P water models such that no solute atom is within 1.0 nm of the box edge. Initial minimisation with

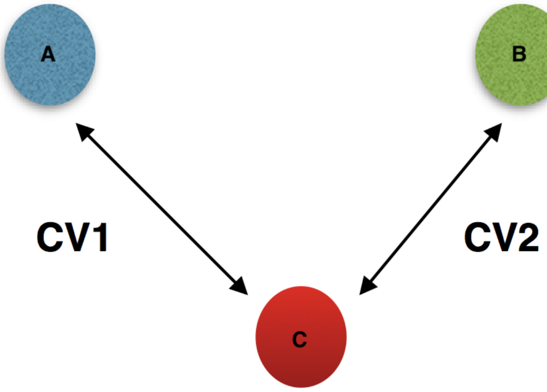


Figure S2: A simplified view of the COM CVs used in this study. **A**, **B** and **C** denotes centre of mass corresponds to flap (residue 58-88), coil (residue 282-302) and catalytic aspartic dyads respectively.

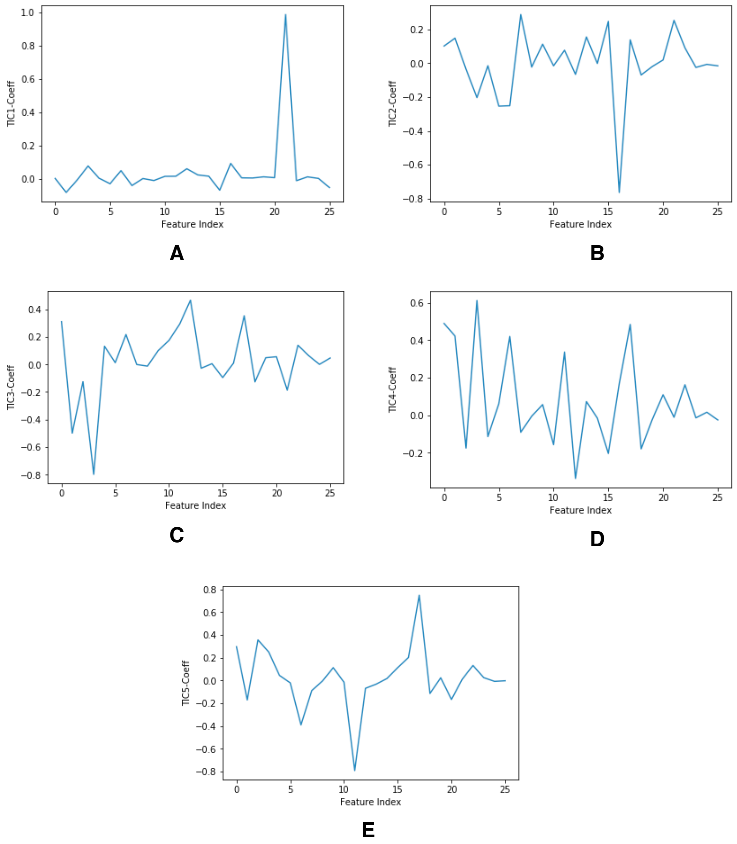


Figure S3: Coefficients corresponding first 5 TICs. The X axis corresponds to the features which are described in Table S3. One can see that TIC1 and TIC3 able to capture rotational degrees of freedom associated with Tyr side-chain. TIC1 gives higher weight to sin$\chi_2$ which makes $\chi_2$ the slowest degrees of freedom.

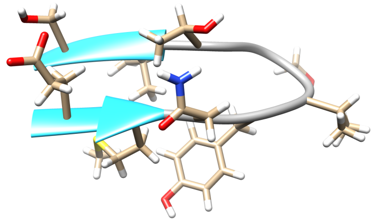

Figure S4: Structural of flap region (residues 73-83) used in generating TICs in case of apo Plm-II.

Table S3: Mean-free sin and cos transformed $\chi_1$ and $\chi_2$ angles and corresponding feature index.

| Feature Index | Name |
|---|---|
| 0 | sin $\chi_1$ 74 |
| 1 | sin $\chi_1$ 75 |
| 2 | sin $\chi_1$ 76 |
| 3 | sin $\chi_1$ 77 |
| 4 | sin $\chi_1$ 78 |
| 5 | sin $\chi_1$ 79 |
| 6 | sin $\chi_1$ 81 |
| 7 | sin $\chi_1$ 82 |
| 8 | sin $\chi_1$ 83 |
| 9 | cos $\chi_1$ 74 |
| 10 | cos $\chi_1$ 75 |
| 11 | cos $\chi_1$ 76 |
| 12 | cos $\chi_1$ 77 |
| 13 | cos $\chi_1$ 78 |
| 14 | cos $\chi_1$ 79 |
| 15 | cos $\chi_1$ 81 |
| 16 | cos $\chi_1$ 82 |
| 17 | cos $\chi_1$ 83 |
| 18 | sin $\chi_2$ 74 |
| 19 | sin $\chi_2$ 75 |
| 20 | sin $\chi_2$ 76 |
| 21 | sin $\chi_2$ 77 |
| 22 | cos $\chi_2$ 78 |
| 23 | cos $\chi_2$ 75 |
| 24 | cos $\chi_2$ 76 |
| 25 | cos $\chi_2$ 77 |

steepest descent algorithm was performed for 5000 steps followed by restrained equilibration (a restrained of 500 kJ/mol was applied on protein $C_\alpha$ and ligand heavy atoms) for 2 ns in the NPT ensemble with Berendsen barostat used to maintain the pressure at 1 bar. The temperature was fixed at 300 K in both cases using the velocity rescaling algorithm. All bonds lengths were constrained using the LINCS algorithm[3] to enable a time-step of 2 fs. A non-bonded cut-off of 1.2 nm was used, and long-range electrostatics were treated by PME using a grid spacing of 0.1 nm. The restrained equilibration was followed by an unrestrained equilibration for 5 ns in the NPT ensemble keeping all other simulation settings identical as before. Finally a snapshot with a volume near the average of this ensemble was used as a starting point for the metadynamics simulation. We have also performed $100ns$ unbiased MD simulation of the protein-ligand complex to check the stability of the ligand in the active site (Figure S23).

In order to estimate the motion necessary for ligand unbinding, we have performed well-tempered metadynamics using the distance between centre of mass of the active site residues[1] and the ligand heavy atoms as CV. This kind of CV was used recently by *Dodda et al.*[4] in order to study unbinding of a small molecule inhibitor from HIV NNRTI. The metadynamics simulation was performed at 300K with a gaussian height of 1.2 kJ/mol and a width at 0.011 nm, deposited every 1 ps. The simulation was stopped when CV distance reached $\sim$ 5 nm.

#### Reweighting with distance

To understand the conformational flexibility of flap and coil region region in apo Plm-II, we devised two distances (DIST2 and DIST3) which quantifies conformational flexibility across all simulations (Figure S5). In case of apo BACE-1, $C_\alpha$ distance between Asp-32 and Thr-72 is referred as DIST2 whereas the $C_\alpha$ distance between Ash-228 and Gln-326 is referred as DIST3. The distances were not used as CVs. In our initial test calculations, we observed that a moving harmonic potential on the distances led to distorted flap conformations. However,

[1]active site residues are: Tyr77, Asp34, Asp214, Ile123, Ser37, Ile32, Val78, Thr217.

we reweighted the free energy landscapes as a function of DIST2 and DIST3 as the distances gave a better quantitative view of the movement of the flap and the coil region. Reweighting the free energy surfaces on DIST2 and DIST3 also gives an understanding of which is the dominant movement between the flap and the coil region. Reweighting was also performed on several H-bond distances involving Tyr.

A reweighting algorithm has been used to calculate the unbiased probability distribution of variables in the well-tempered metadynamics simulations.[5] If a bias is acting on a system, it is constantly changing the biased probability distribution of the system. Using the reweighting algorithm, we can remove the effect of bias and recover the unbiased probability distribution along chosen degrees of freedom.

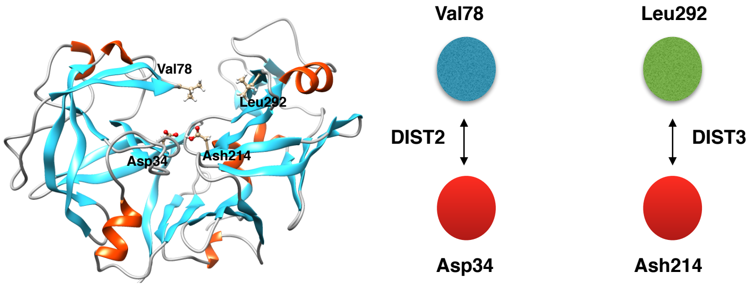


Figure S5: Representation of distance matrices, DIST2 and DIST3 used in this study to quantify movement of flap (DIST2) and coil region (DIST3) in apo Plm-II.

#### Computational tools

All simulations were performed using *Gromacs 2018*[6] patched with *Plumed*.[7] The *tLeap* module of Amber was used to generate topology and co-ordinate files. *Acpype*[8] was used to convert the Amber topology and co-ordinates to a *Gromacs* compatible version.

### Normal state

The normal state can be divided into two parts: $N-$ ($-\frac{\pi}{3}$ $radian$) and $N+$ ($\frac{\pi}{3}$ $radian$). $N+$ conformation is unstable because the $\gamma$ atom ($CG$) of tyrosine is in close contact with backbone $CO$ group (Figure S6). In BACE-1, H-bond to Lys was formed due to $N+$ orientation of tyrosine.

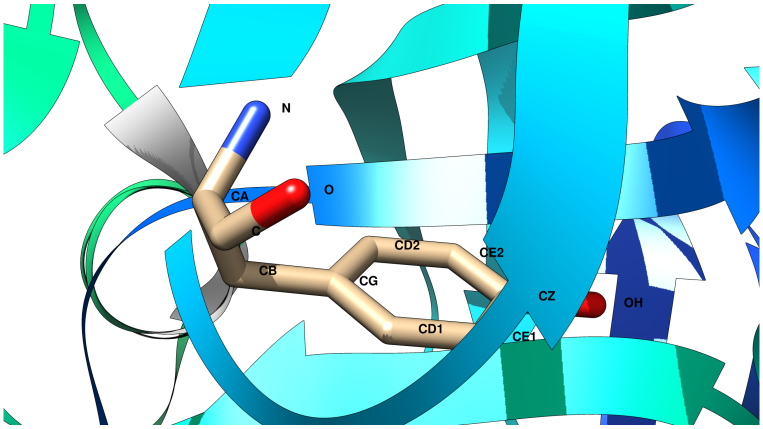


Figure S6: $N+$ conformation of tyrosine.

#### Clustering

Clustering was performed using TIP3P MD simulation (denoted as R1-T3P) of apo Plm II. We used *K-centers* clustering algorithm integrated with Python package enspara[9] (`https://github.com/bowman-lab/enspara`) to generate 10000 cluster centers. Clusters were defined geometrically using backbone RMSD of residues 73-88. For each cluster centers we have calculated DIST2, DIST3, distance between Tyr–Trp and Tyr–Asp using *Plumed*. The scripts can be accessed here: `https://github.com/sbhakat/Plasmepsin-bace/tree/master/Clustering-script`. Figure S20 and S22 compares some of the representative cluster centers with crystal conformation of apo Plm-II (PDB: 1LF4).

### Statistical analysis

We have calculated time dependent free energy difference between flipped and normal ($-\frac{\pi}{3}$ *radian*) state in Torsion-Metad simulations. We integrated multiple free-energy profiles in two basins defined by the following intervals on $\chi_1$ space: flipped ($-2.5 < \chi_1 < -3.14$);

normal ($-1.5 < \chi_1 < -0.5$). We have also calculated free energy difference between H-bond to Trp and Asp and different basins along DIST2 (Figure S6, S7 and S8).

Free energy difference was also calculated between $N-$ ($-\frac{\pi}{3}$ *radian* along $\chi_1$) and $N+$ ($\frac{\pi}{3}$ *radian* along $\chi_1$) conformations of the normal state.

The block averages are calculated using reweighted free energy surface along $\chi_1$ over a period of 50 ns and the precision is approximated by standard deviation in each block.

Statistical analyses was performed using a bootstrapping procedure with 100 repeats.

Table S4: Block averaging performed on the free energy difference between flipped and normal state in two independent Torsion-Metad simulations with TIP3P water model.

| | Average (kJ/mol) |
|---|---|
| **FF14SB-R1** | |
| 600-650 ns | 1.24±0.004 |
| 650-700 ns | 1.11±0.006 |
| 700-750 ns | 0.98±0.005 |
| 750-800 ns | 0.86±0.004 |
| 800-850 ns | 0.75±0.004 |
| **FF14SB-R2** | |
| 600-650 ns | -0.05±0.02 |
| 650-700 ns | -0.05±0.02 |
| 700-750 ns | -0.05±0.01 |

Table S5: Block averaging performed on the free energy difference between flipped and normal state in Torsion-Metad simulation starting with SN conformation.

| Interval (ns) | Average (kJ/mol) |
|---|---|
| 300-350 | -2.03±0.02 |
| 350-400 | -2.03±0.02 |
| 400-450 | -2.03±0.02 |
| 450-500 | -2.03±0.02 |
| 500-550 | -2.02±0.02 |

Table S6: Free energy difference in metadynamics simulations using apo Plm-II. $Trp - Asp$ defines free energy difference between H-bond to Trp and Asp. It was calculated using interval of $0.27 - 0.35$ nm. and $0.65 - 0.90$ nm along 1D reweighted free energy profile along H-bond to Trp. Free energy difference along DIST2 was calculated using an interval of $1.00 - 1.15$ nm and $1.15 - 1.6$ nm. N $\pm$ defines the free energy difference along $\chi_1$ using the interval $-1.4 - -0.8$ radian and $0.8 - 1.9$ radian. The time average was performed using last part of the simulations (500 ns onwards). Free energy values are in kJ/mol.

| System | $Flipped - normal$ | $Trp - Asp$ | DIST2 | N± |
|---|---|---|---|---|
| FF14SB-R1 | 1.02±0.01 | -0.79±0.01 | 0.82±0.01 | -10.49±0.13 |
| FF14SB-R2 | -0.61±0.01 | 0.71±0.01 | -0.09±0.01 | -8.90±0.03 |
| TIP4P-Ew | 0.62±0.01 | -0.74±0.02 | -0.35±0.01 | -7.32±0.01 |
| CHARMM | -3.16±0.01 | 3.45±0.02 | -3.18±0.01 | -8.35±0.01 |

Table S7: Free energy difference of flap opening was calculated using 1D free energy surface reweighted on DIST2 using an interval of $1.00 - 1.2$ nm and $1.8 - 2.2$ nm. Free energy values are in kJ/mol.

| System | Open |
|---|---|
| FF14SB-R1 | -2.78±0.01 |
| FF14SB-R2 | -2.71±0.02 |
| TIP4P-Ew | -3.08±0.02 |
| CHARMM | -5.53±0.01 |

Table S8: Free energy difference in metadynamics simulations starting with SO, SN and SSO conformations. $Trp - Asp$ defines free energy difference between H-bond to Trp and Asp. It was calculated using interval of $0.27 - 0.35$ nm and $0.65 - 0.93$ nm along 1D reweighted free energy profile along H-bond to Trp. Free energy difference along DIST2 was calculated using an interval of $1.00 - 1.15$ nm and $1.15 - 1.4$ nm. N $\pm$ defines the free energy difference along $\chi_1$ using the interval $-1.4 - -0.8$ radian and $0.8 - 1.9$ radian. The time average was performed using last part of the simulations (300 ns onwards). Free energy values are in kJ/mol.

| System | $Flipped - normal$ | $Trp - Asp$ | DIST2 | N± |
|---|---|---|---|---|
| SSO | -3.53±0.02 | 3.17±0.02 | -4.09±0.02 | -6.51±0.03 |
| SO | -1.68±0.01 | 1.22±0.01 | -0.83±0.01 | -14.12±0.03 |
| SN | -2.10±0.01 | 1.57±0.01 | -1.61±0.01 | -13.67±0.02 |

Table S9: Free energy difference between H-bond to Lys and Trp was calculated using reweighted 1D free energy surface along H-bond to Lys using intervals $0.25 - 0.38$ and $0.9 - 1.2$ nm. Free energy difference between H-bond to Ser and Trp was calculated using reweighted free energy surface along H-bond to Ser using intervals $0.25-0.38$ and $0.38-0.57$ nm. Free energy values are in kJ/mol.

| System | Hbond Lys | Hbond Ser |
|---|---|---|
| SSO | 13.50±0.02 | 14.89±0.01 |
| SO | 18.83±0.02 | 11.33±0.04 |
| SN | 19.17±0.02 | 7.60±0.1 |

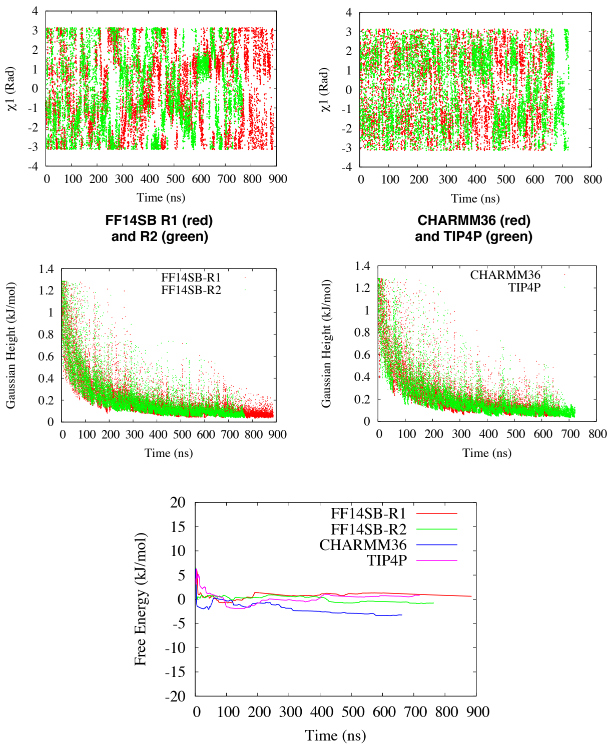


Figure S7: Convergence behavior of Torsion-Metad simulations of Plm-II. The upper panel shows the fluctuation of the $\chi_1$ angle in various metadynamics simulations. The middle panel shows the Gaussian height (kJ/mol) as a function of simulation time for the same simulations. The lower panel shows the estimated free energy between the flipped and normal states ($-\frac{\pi}{3}$ rad) as a function of simulation time.

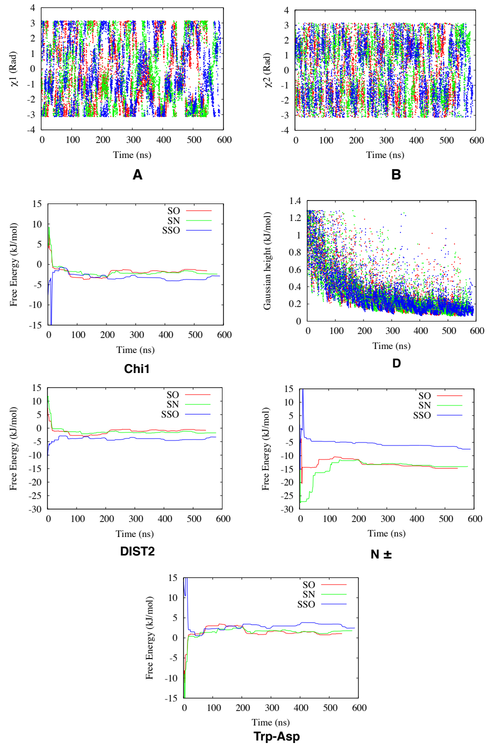


Figure S8: Convergence behavior of Torsion-Metad simulations of BACE-1. The diagrams show the fluctuation of $\chi_1$ (A) and $\chi_2$ (B) starting from the SO (red), SN (green) and SSO (blue) structures, respectively, and the Gaussian height as a function of simulation time (D). The free-energy difference between the flipped and normal states estimated along $\chi_1$ (C), DIST2 (E), $N\pm$ (F; described in the Supplementary Information) and $Tyr - Asp$ (G) as a function of simulation time is also shown. The intervals defining the states are shown in Table S8 in the Supplementary Information.

Table S10: Sampling of open conformations (DIST2 > 1.8 nm.) in MD (TIP3P water model) and metadynamics simulations. Tor-Metad-1 and Tor-Metad-2 correspond to two independent metadynamics runs with torsion CVs.

| Simulation | % Open |
|---|---|
| R1-T3P | 8 |
| R2-T3P | 0.2 |
| R3-T3P | 0.2 |
| R4-T3P | 0.4 |
| Tor-Metad-1 | 12 |
| Tor-Metad-2 | 9 |

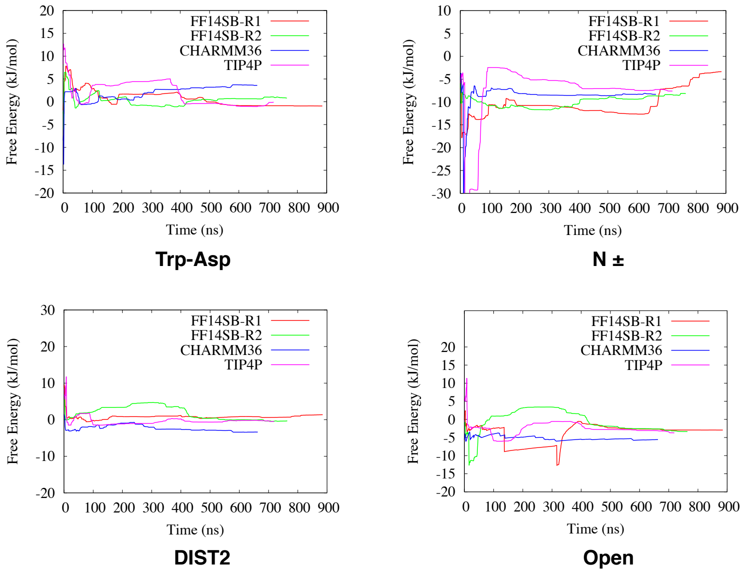


Figure S9: Free energy difference along different collective variables in metadynamics simulations with apo Plm-II. The intervals were mentioned in Table S6 and S7

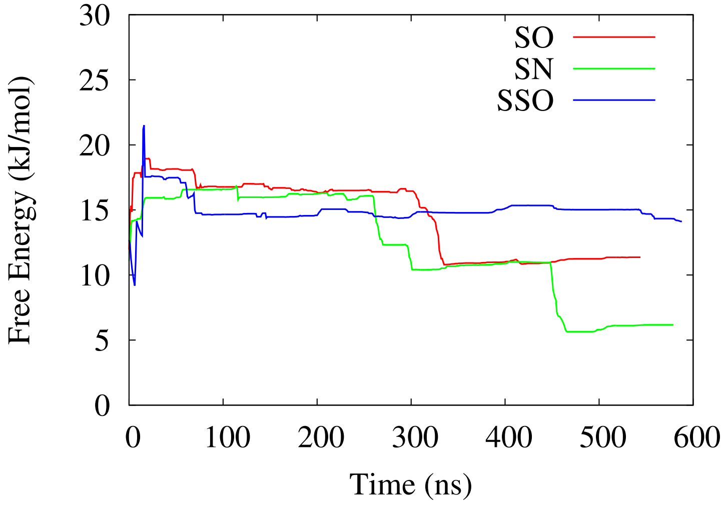


Figure S10: Free energy difference between H-bond to Ser and Trp as a function of simulation time for apo BACE-1.

Table S11: Percent hydrogen bond population in two independent MD runs (R1 and R2) of apo Plm II with TIP3P water molecules.

| Simulation | Tyr-Trp | Tyr-Asn | Tyr-Ser | Tyr-Asp | Tyr-Gly |
|---|---|---|---|---|---|
| R1-T3P | 61 | 0.2 | 10 | 0 | 0 |
| R2-T3P | 40 | 7 | 31 | 0 | 0 |

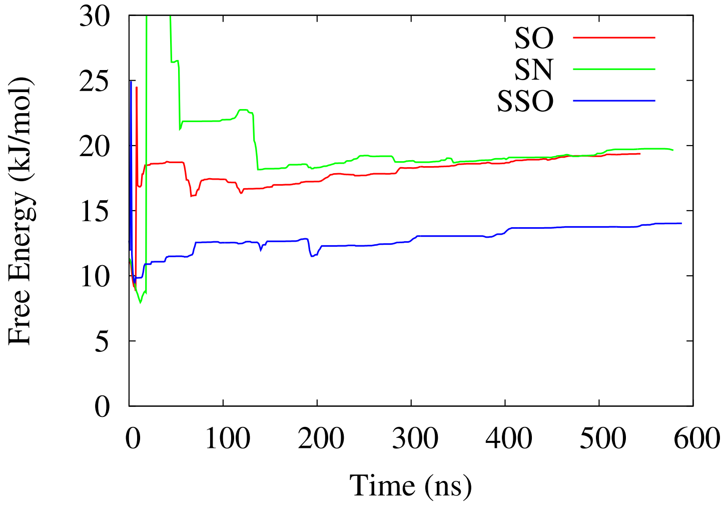


Figure S11: Free energy difference between H-bond to Lys and Trp as a function of simulation time for apo BACE-1.

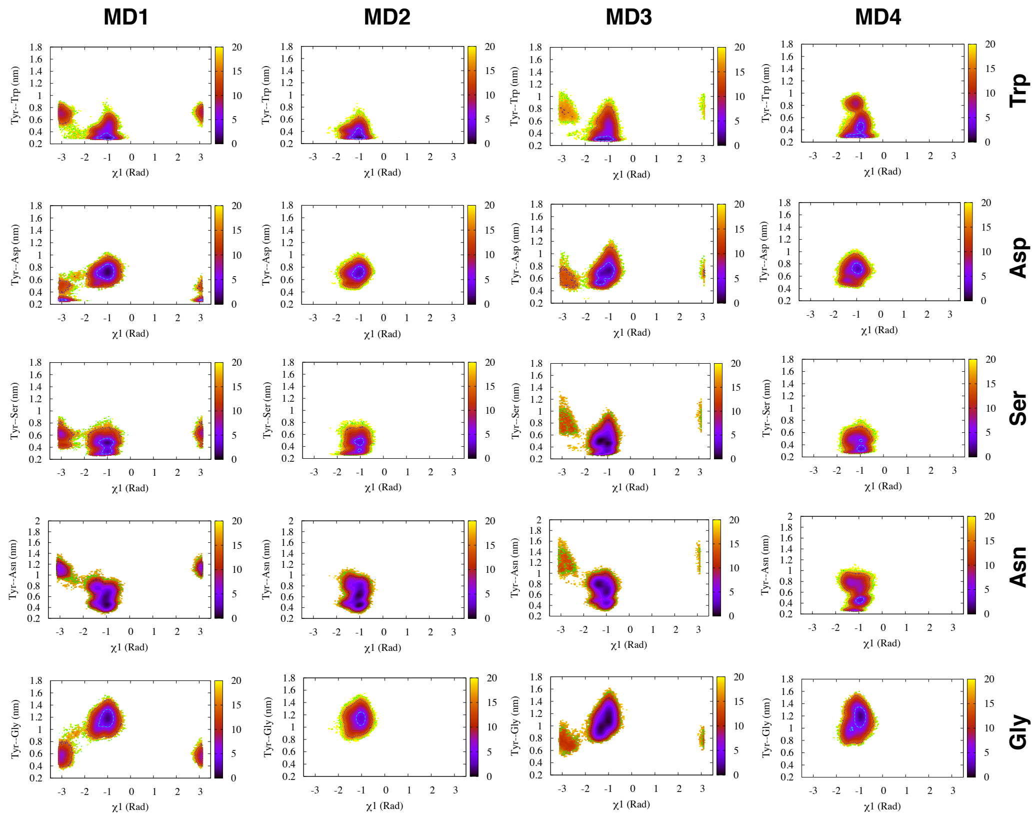


Figure S12: Free energy surface projected on $\chi_1$ and H-bond distances in case of independent MD simulations with TIP3P water model in apo Plm II. **MD1** did a slight sampling of normal and flipped states. The flipped state was stabilised by formation of H-bond to Asp. In case of **MD1-MD4**, the normal state was stabilised by inter-changing H-bonds to Trp and Ser. **MD4** sampled additional H-bond to Asn.

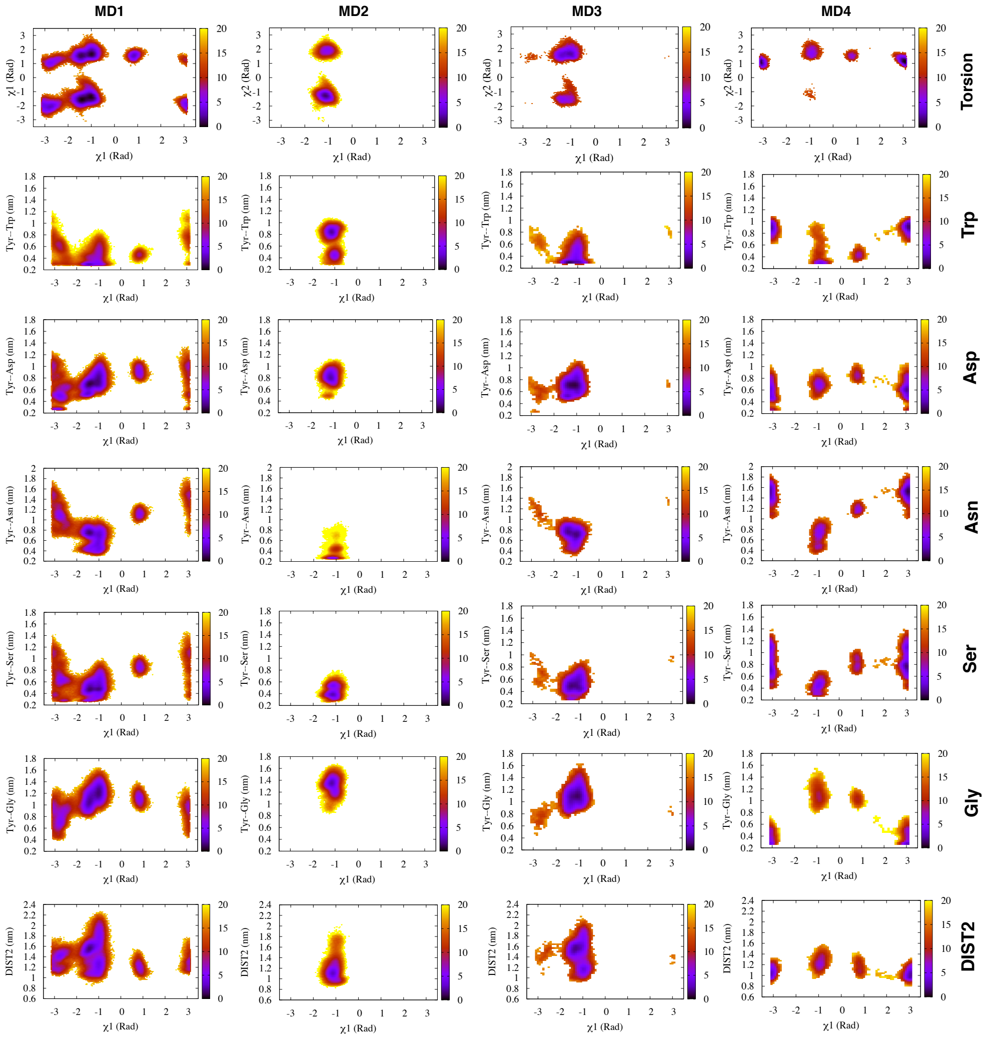


Figure S13: Free energy surface projected on $\chi_1$ and other variables (e.g. $\chi_2$, H-bond distances (Trp, Asp, Ser, Asn and Gly) and DIST2) in case of four independent MD simulations with TIP4P-Ew water-model in apo Plm II. **MD1** and **MD4** sampled both normal and flipped states. The flipped state in **MD4** was stabilised by inter-changing H-bond to Asp and Gly.

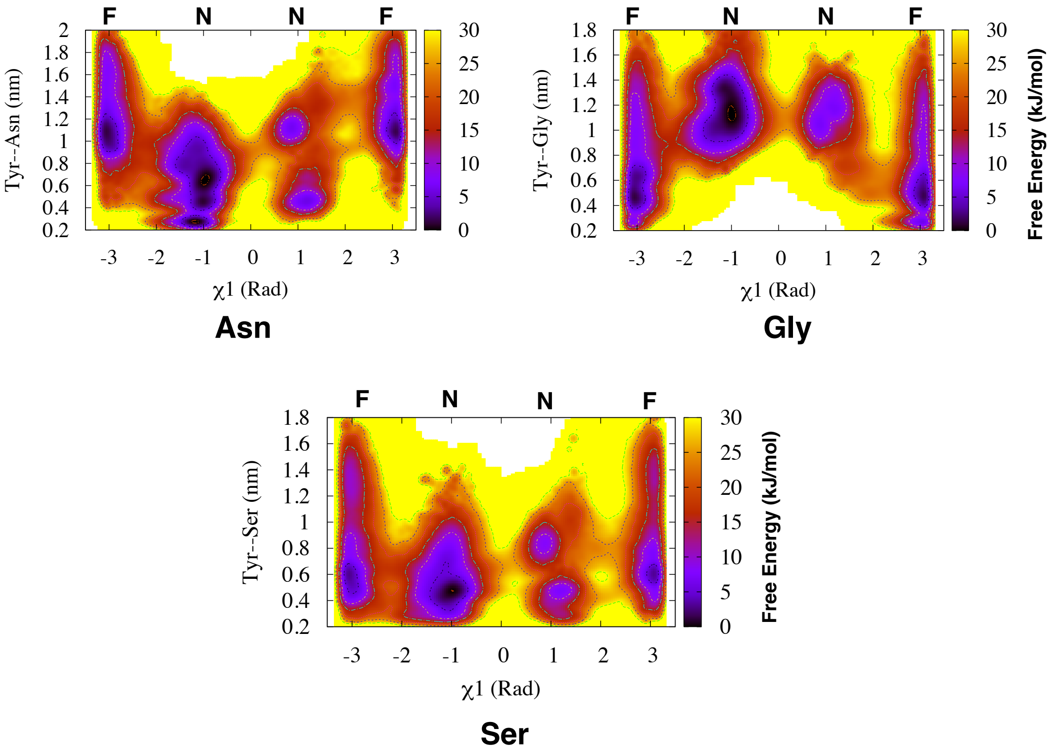


Figure S14: Free energy surface reweighted on $\chi_1$ and H-bond distances (Asn, Ser and Gly) in case of Torsion-MetaD simulations with apo Plm-II. **F** and **N** denotes flipped and normal states respectively.

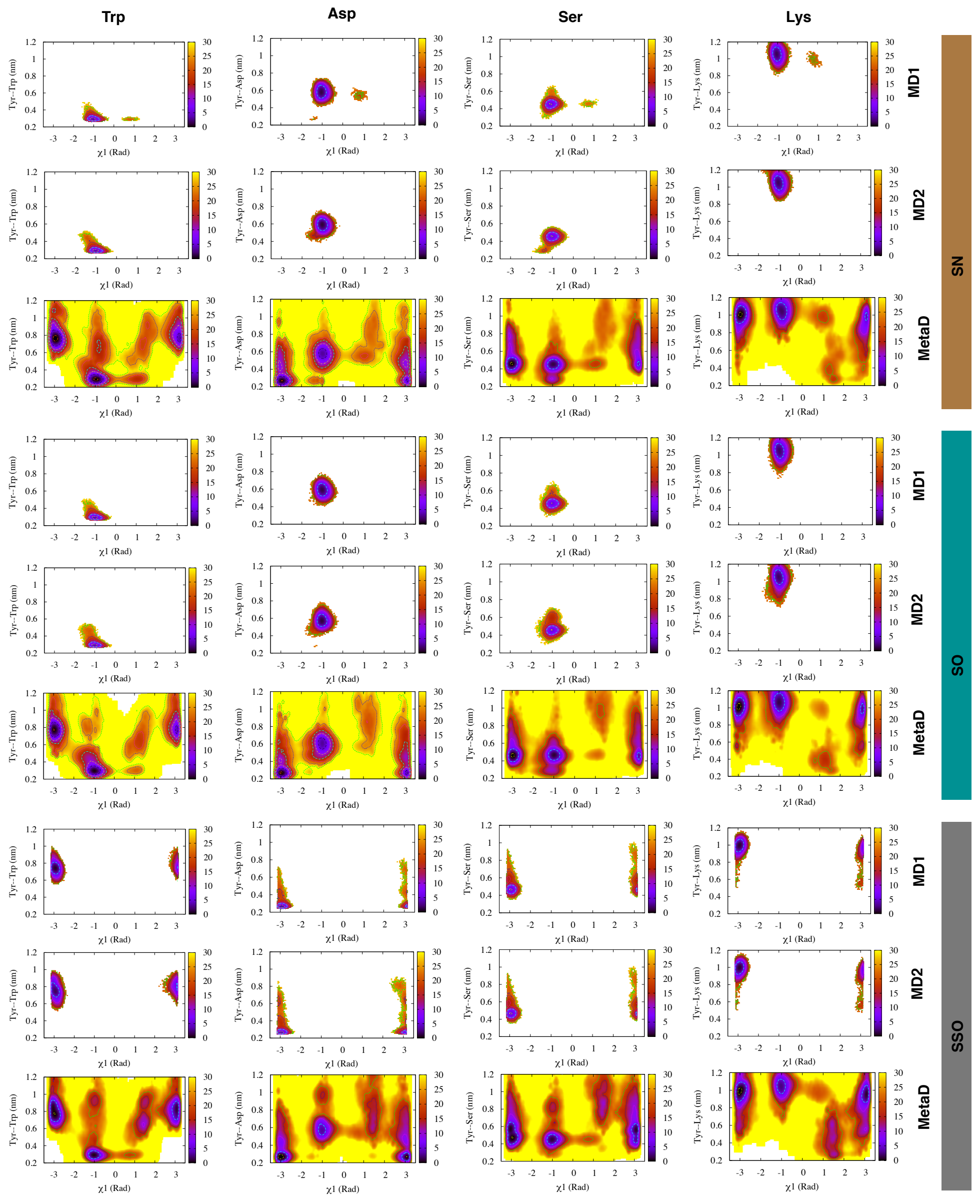


Figure S15: Free energy surface projected on $\chi_1$ and H-bond distances in case of MD and Torsion-Metad simulations starting with SN, SO and SSO conformations. MD simulations starting with SO and SN conformations only sampled the normal state. Whereas, simulations starting with SSO conformation remained stuck in the flipped state. Torsion-Metad starting with SO, SN and SSO conformations sampled both normal and flipped states.

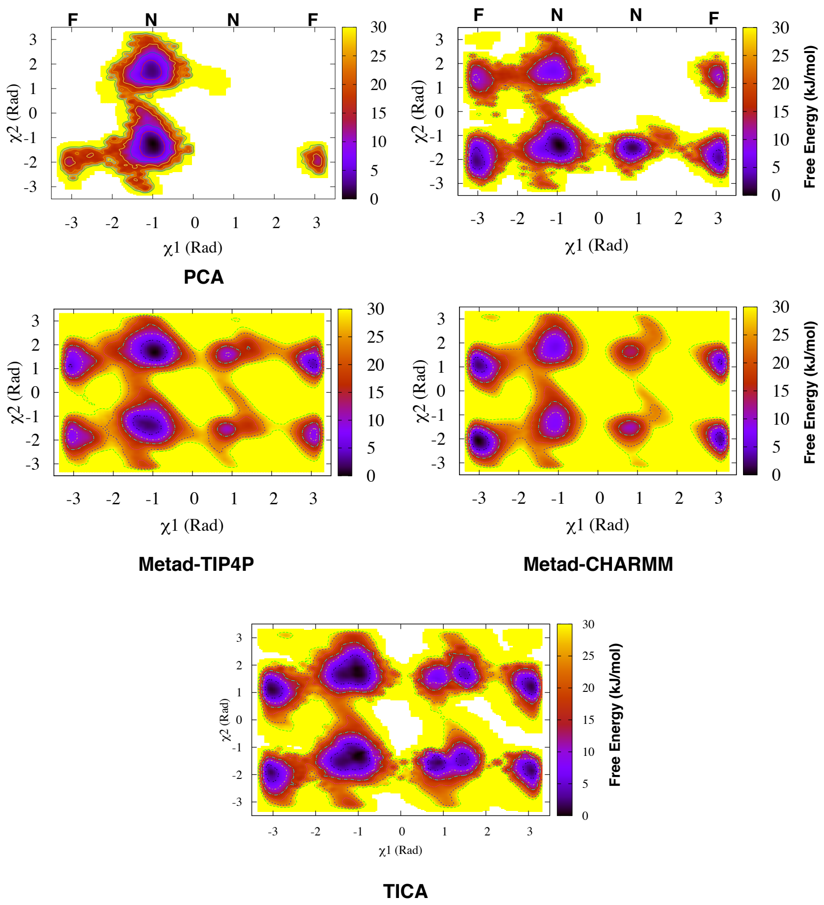


Figure S16: Free energy surface reweighted on $\chi_1$ and $\chi_2$ for metadynamics simulations using PCA, COM, torsion (denoted as Metad-TIP4P and Metad-CHARMM), TIC CVs.

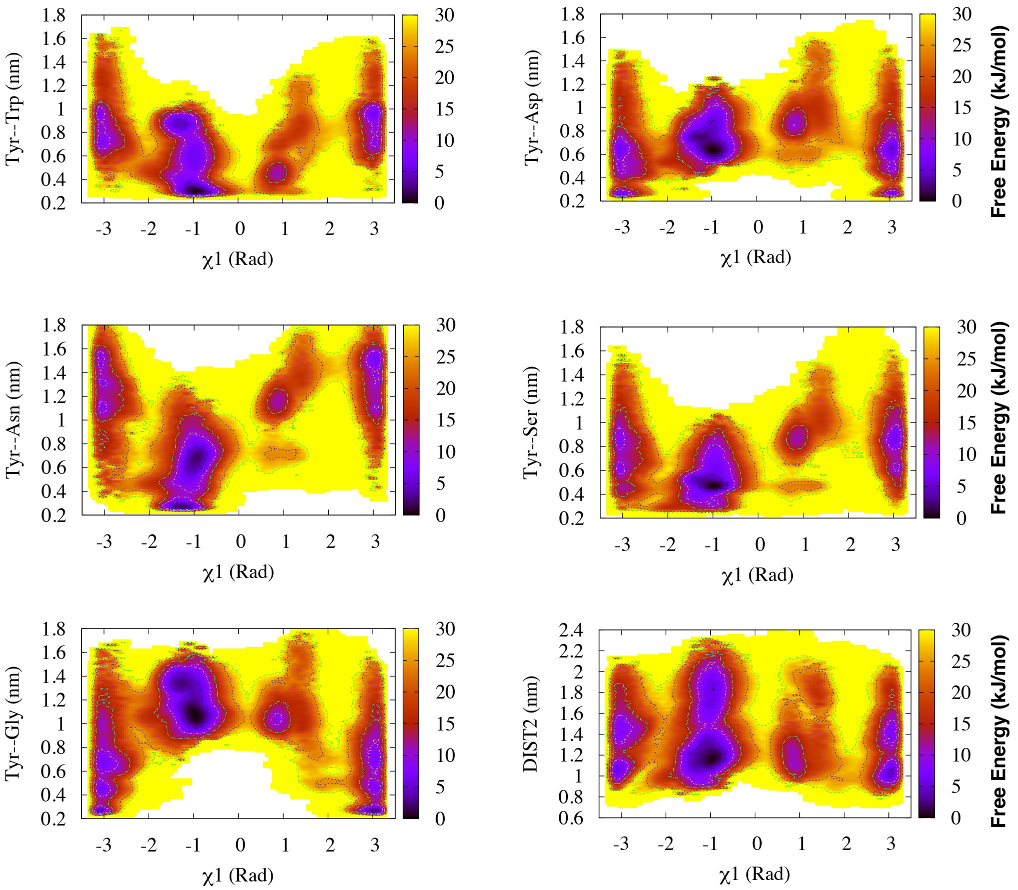


Figure S17: 2D free energy surface reweighted on $\chi_1$ and distances (H-bond to Trp, Asp, Ser, Asn, Gly and DIST2) in Torsion-Metad simulation with TIP4P-Ew water model in apo Plm-II.

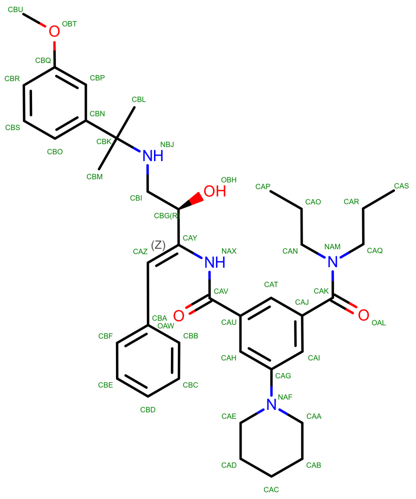


Figure S18: Structure of the ligand bound with PDB: 4Y6M `https://www.rcsb.org/structure/4Y6M`.

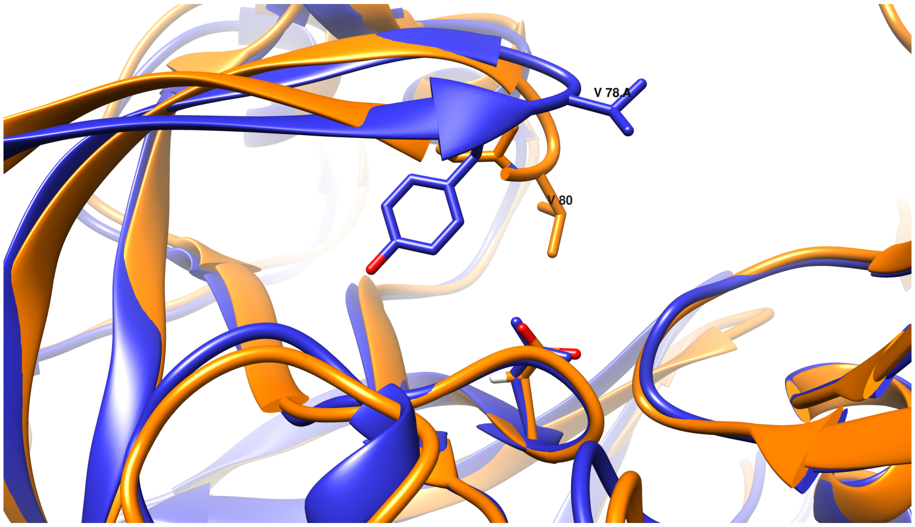


Figure S19: Structural representation of collapsed flap (Tyr to Ala mutation) conformation (ORANGE) with respect to apo Plm-II (PDB: 1LF4) (BLUE). The orientation of Val with respect to catalytic Asp gives a structural insight of the flap collapse.

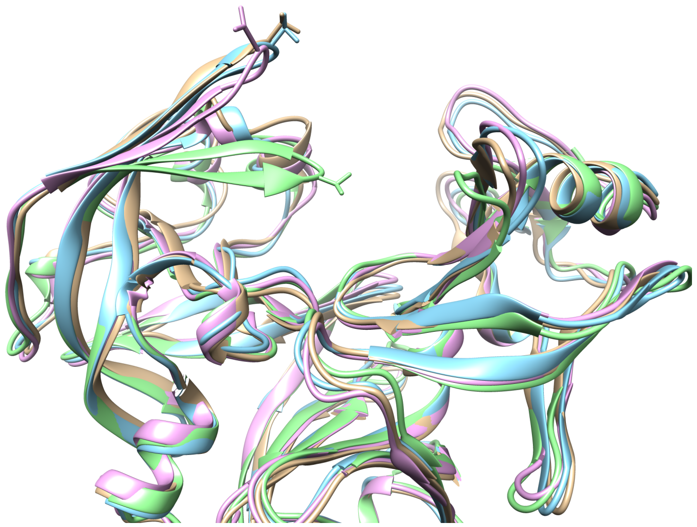

Figure S20: Snapshot of wide open conformations (DIST2>2.3 nm) generated from clustering with relative to apo Plm-II (PDB:1LF4, highlighted in green). The flap tip valine residues are highlighted for comparison.

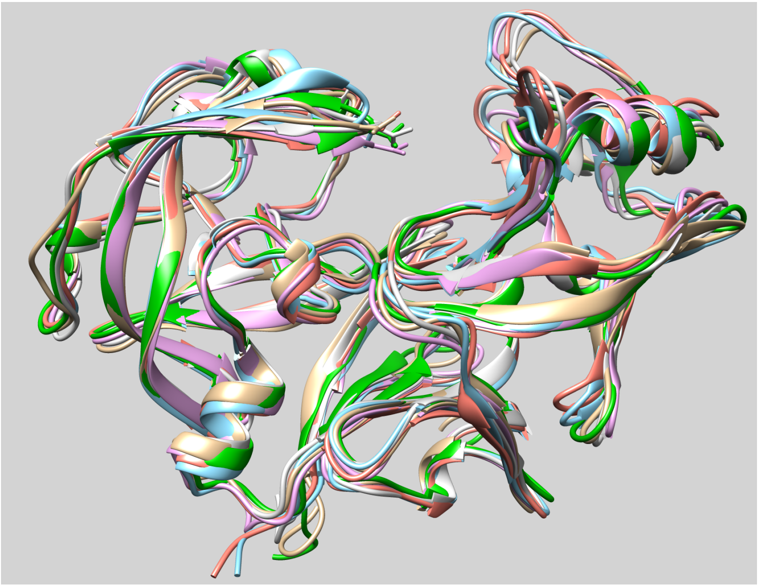

Figure S21: Snapshot of conformations with DIST2∼1.2 nm generated from clustering with relative to apo Plm-II (PDB:1LF4, highlighted in green). The flap tip valine residues are highlighted for comparison.

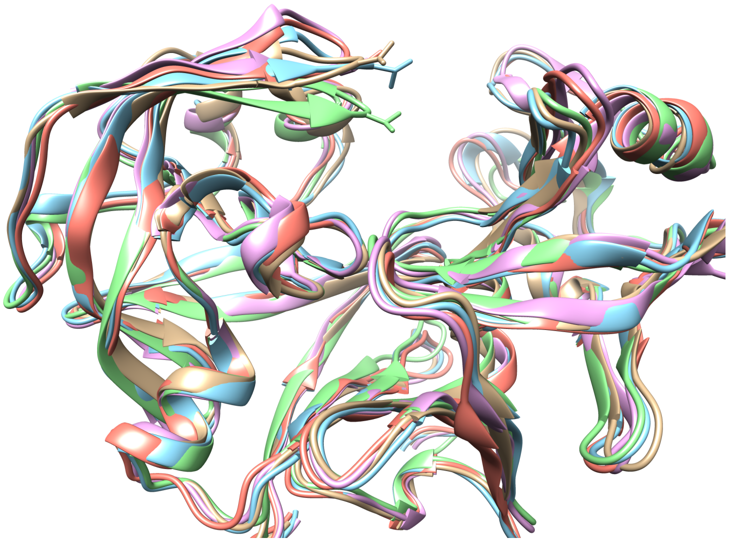

Figure S22: Snapshot of conformations with DIST2∼1.6 nm generated from clustering with relative to apo Plm-II (PDB:1LF4, highlighted in green). The flap tip valine residues are highlighted for comparison.

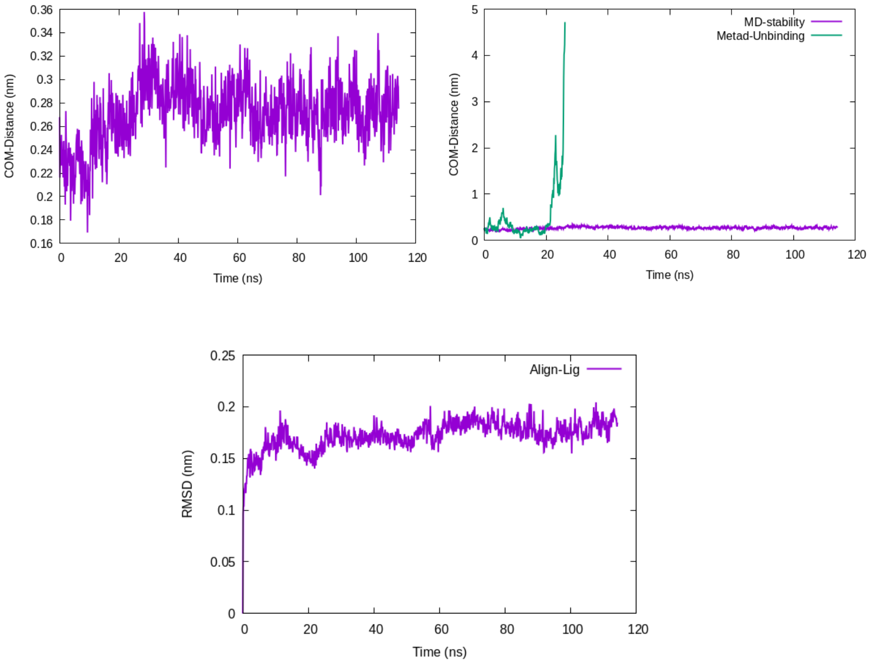


Figure S23: Equilibrium fluctuation of COM-Distance in MD simulation of protein ligand complex (upper left panel) and its comparison with metadynamics based unbinding simulation (upper right panel). RMSD of ligand in the active site shows the stability of the ligand during MD simulation (bottom panel). RMSD of the ligand was calculated by aligning on the ligand heavy atoms.

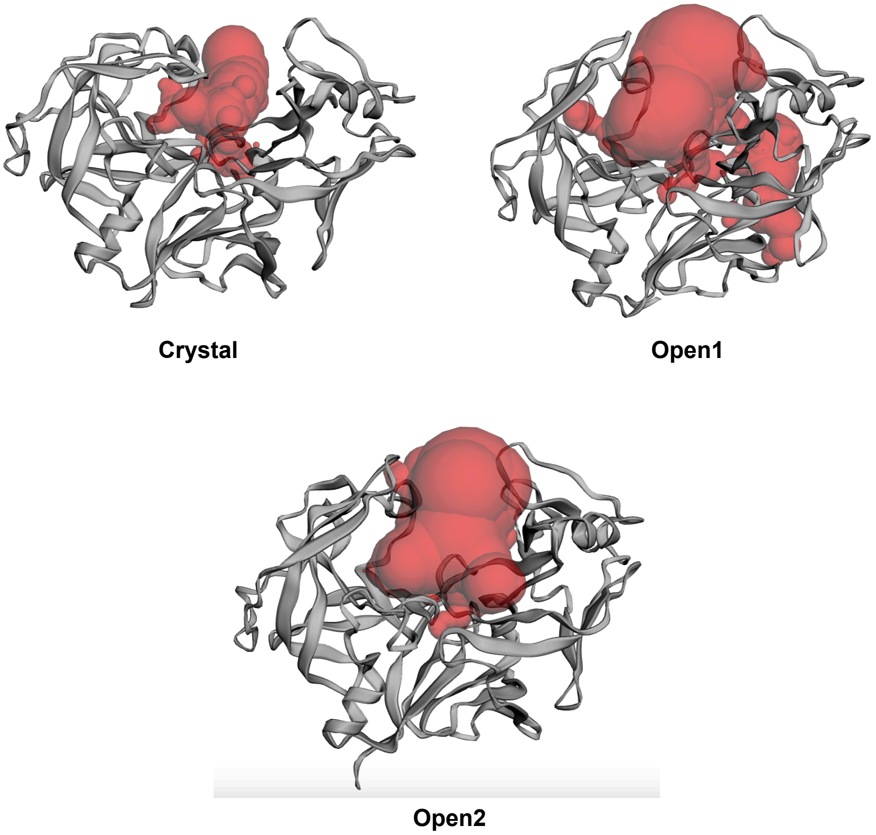


Figure S24: Pocket prediction using CASTp server.[10] For equilibrated crystal conformation the pocket volume is predicted to be 692.27 $\AA^3$. Whereas for $Open1$ and $Open2$ conformations $(DIST2 > 2.3nm)$ the pocket surface areas are predicted to be 2056.63 and 2534.25 $\AA^3$. respectively.

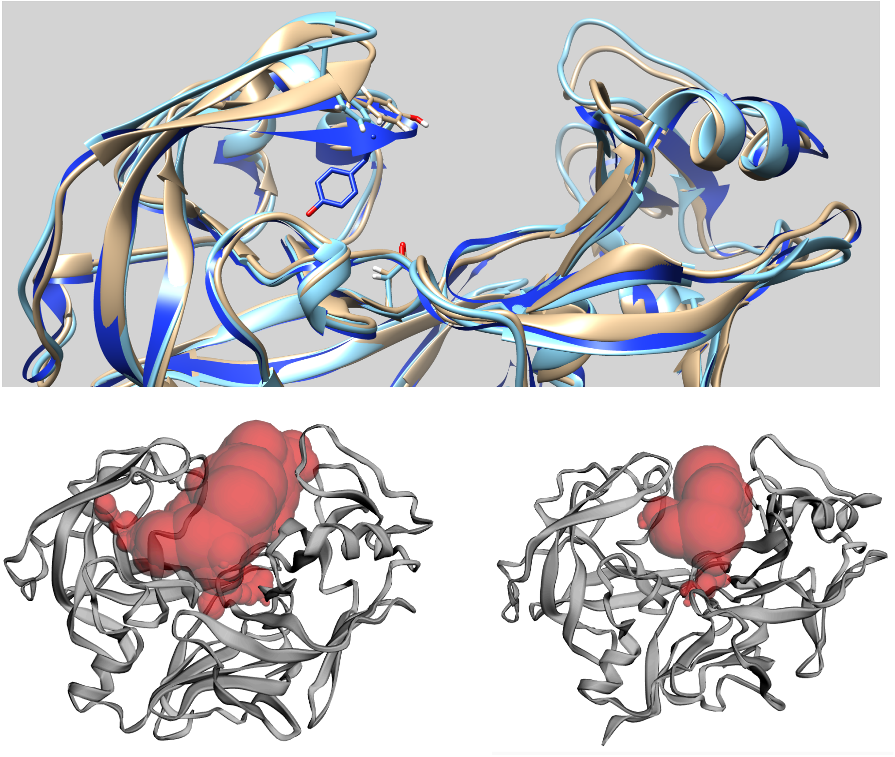

Figure S25: Plm II flap in open conformations ($DIST2 > 1.8nm$) with Tyr in flipped conformation (apo Plm II, PDB:1LF4 has been depicted in BLUE for reference). Pocket prediction using CASTp server[10] shows pocket volume of 1706.72 (Open1-flipped) and 893.9 (Open2-flipped) $\AA^3$ respectively for snapshots corresponding these two conformations.
